# High-Affinity, Structure-Validated and Selective Macrocyclic Peptide Tools for Chemical Biology Studies of Huntingtin

**DOI:** 10.1101/2025.08.06.668955

**Authors:** Esther Wolf, Rebeka Fanti, Tatsuya Ikenoue, Justin C. Deme, Swati Balakrishnan, Brandon A. Keith, Matthew G. Alteen, Renu Chandrasekaran, Manisha Yadav, Ritika Bhajiawala, Suzanne Ackloo, Jia Feng, Mahmoud A. Pouladi, Aled M. Edwards, Derek Wilson, Susan M. Lea, Hiroaki Suga, Rachel J. Harding

**Affiliations:** Structural Genomics Consortium, University of Toronto, Toronto, ON M5G 1L7, Canada; Department of Molecular Genetics, Faculty of Medicine, University of Toronto, Toronto, ON M5S 1A8, Canada; Department of Chemistry, Graduate School of Science, The University of Tokyo, Tokyo 113-0033, Japan; Center for Structural Biology, Center for Cancer Research, National Cancer Institute, Frederick, MD, USA; Department of Medical Biophysics, University of Toronto, Toronto, ON, M5G1L7, Canada; Department of Chemistry, York University, Toronto, ON M3J 1P3, Canada; Department of Medical Genetics, Centre for Molecular Medicine and Therapeutics, Djavad Mowafaghian Centre for Brain Health, Edwin S. H. Leong Centre for Healthy Aging, Faculty of Medicine, University of British Columbia, British Columbia Children’s Hospital Research Institute, Vancouver, BC, Canada; Leslie Dan Faculty of Pharmacy, University of Toronto, Toronto ON M5S 3M2, Canada; Department of Pharmacology and Toxicology, Faculty of Medicine, University of Toronto, Toronto, ON M5S 1A8, Canada

**Keywords:** Huntingtin, HAP40, peptide macrocycles, Huntington’s disease, ligand discovery

## Abstract

Huntington’s disease (HD) is a fatal neurodegenerative disorder caused by a CAG repeat expansion in the *Huntingtin* (*HTT*) gene, with no disease-modifying therapies currently available. The precise molecular function of the HTT protein is unclear, and the lack of selective chemical tools has limited functional studies. We have identified and characterized macrocyclic peptide binders targeting HTT. These binders exhibit low-nanomolar affinity *in vitro* and engage distinct HTT and HTT-HAP40 interfaces, as revealed by hydrogen-deuterium exchange mass spectrometry and cryo-electron microscopy. Chemoproteomics confirmed selective binding in cell extracts from wildtype but not HTT-null cell lines. HAP40 consistently and stoichiometrically co-purified with HTT across cell lines, including with HTT variants containing different CAG repeat lengths, highlighting the broad presence of the HTT-HAP40 complex.

**Significance Statement:** Huntingtin (HTT) is a large, essential protein with conserved roles in development, intracellular trafficking, and protein homeostasis, yet its precise molecular functions remain incompletely defined. Here, we report the first high-affinity, selective, and structurally validated macrocyclic peptide ligands for HTT. These chemical tools bind HTT and its complex with HAP40 across polyglutamine repeat lengths, enabling direct interrogation of HTT structure and function in health and disease contexts. By overcoming longstanding barriers to studying HTT at the molecular level, these ligands open new avenues for discovery in neurobiology, cell biology, and offer opportunities for therapeutic development. This study delivers urgently needed tools to both the Huntington’s disease field and the broader scientific community seeking to understand this elusive and biologically fundamental protein.

## Introduction

Huntington’s disease (HD) is a rare, debilitating autosomal dominant neurodegenerative disorder caused by expansion of a CAG repeat tract in the *Huntingtin* (*HTT)* gene (MacDonald et al., 1993) above 35 repeats (Shelbourne et al., 2007). Symptoms typically begin in adulthood with a life expectancy of ~15 years after clinical diagnosis (Ross & Tabrizi, 2011). HD manifests as cognitive, psychiatric and movement symptoms primarily attributed to progressive degeneration of medium spiny neurons in the striatum (Aylward et al., 2011). Despite the identification of this genetic cause of HD over three decades ago, effective disease-modifying treatments remain elusive (Tabrizi et al., 2022).

HTT is a large 3144 amino acid protein organized into N- and C-terminal globular domains of HEAT (huntingtin, elongation factor 3, protein phosphatase 2a, and yeast kinase TOR1) repeats(Saudou & Humbert, 2016). The CAG-repeat encoded polyglutamine (polyQ) tract, which is encoded by exon 1 of the *HTT* gene, is not resolved in cryo electron microscopy (cryo-EM) structures of the full-length protein (Guo et al., 2018). PolyQ expanded protein (mHTT) has high structural similarity to wildtype HTT (Huang et al., 2021) and is predicted to differ mostly in the conformational space accessed by the flexible polyglutamine expanded exon 1 region (Harding et al., 2021a), reviewed by Seefelder et al.

HTT is ubiquitously expressed and has been implicated in a variety of cellular processes, including autophagy (Rui et al., 2015), apoptosis (Luo & Rubinsztein, 2009; Raychaudhuri et al., 2008), mitosis (Godin et al., 2010), neuronal signaling (Tang et al., 2003), vesicular trafficking (Pardo et al., 2010), and the endosomal pathway (Pal et al., 2006). HTT has been hypothesized to serve as a scaffold for protein-protein (Greco et al., 2022; Maiuri et al., 2017; Ratovitski et al., 2012) and protein-RNA interactions (Culver et al., 2016; Yadav et al., 2024). However, 40-kDa huntingtin-associated protein (HAP40) (Peters & Ross, 2001) remains the only structurally characterized interactor in the published literature of the hundreds reported (Aaronson et al., 2021). HTT forms an extensive interaction interface with HAP40 which confers structural stability to HTT (Seefelder et al., 2022).

The precise mechanisms by which HTT exerts its putative scaffolding function require further investigation, including its HAP40 dependent and independent partners, subcellular localization, and the impact of polyQ expansion. Uncovering these mechanisms will be enabled by high-quality research tools, including antibodies and chemical tools. Previously, we characterized tens of commercially available antibodies in a range of cell biology studies to identify those that were most potent and selective in several applications (Fanti et al., 2024). Here, we discover and characterize potent and selective macrocycle chemical tools for HTT and demonstrate their utility by using them to characterize the HTT protein network, identifying HTT-HAP40 as the dominant endogenous proteoform.

## Results

### Discovery of macrocycles targeting HTT and HTT-HAP40

Full-length wildtype HTT with a polyQ tract of 23 glutamines in either its apo form (HTTQ23) or HAP40 bound form (HTTQ23-HAP40) was purified (Alteen et al., 2023; Harding et al., 2019; Hutchinson & Seitova, 2021) and deployed into the RaPID platform coupled to Flexible *in vitro* Translation (FIT) (Fig. 1b) (Goto et al., 2011; Murakami et al., 2006). Briefly, FIT harnesses custom charged tRNAs and *in vitro* reconstituted translation to repurpose traditional codons (e.g., AUG) and generate peptides with nonstandard residues. In place of methionine, the reaction mixture contains N-chloroacetylated tyrosine (ClAc-Tyr) charged tRNA (AUG anticodon). Therefore, spontaneous cyclization is enabled via S_N_2 nucleophilic attack of initiator ClAc-Tyr by a downstream cysteine, forming an irreversible thioether linkage (Goto & Suga, 2021) (Fig. 1c). The translation reaction is then terminated using puromycin, linking each macrocycle to its cognate mRNA (i.e., mRNA display).

**Figure 1.**
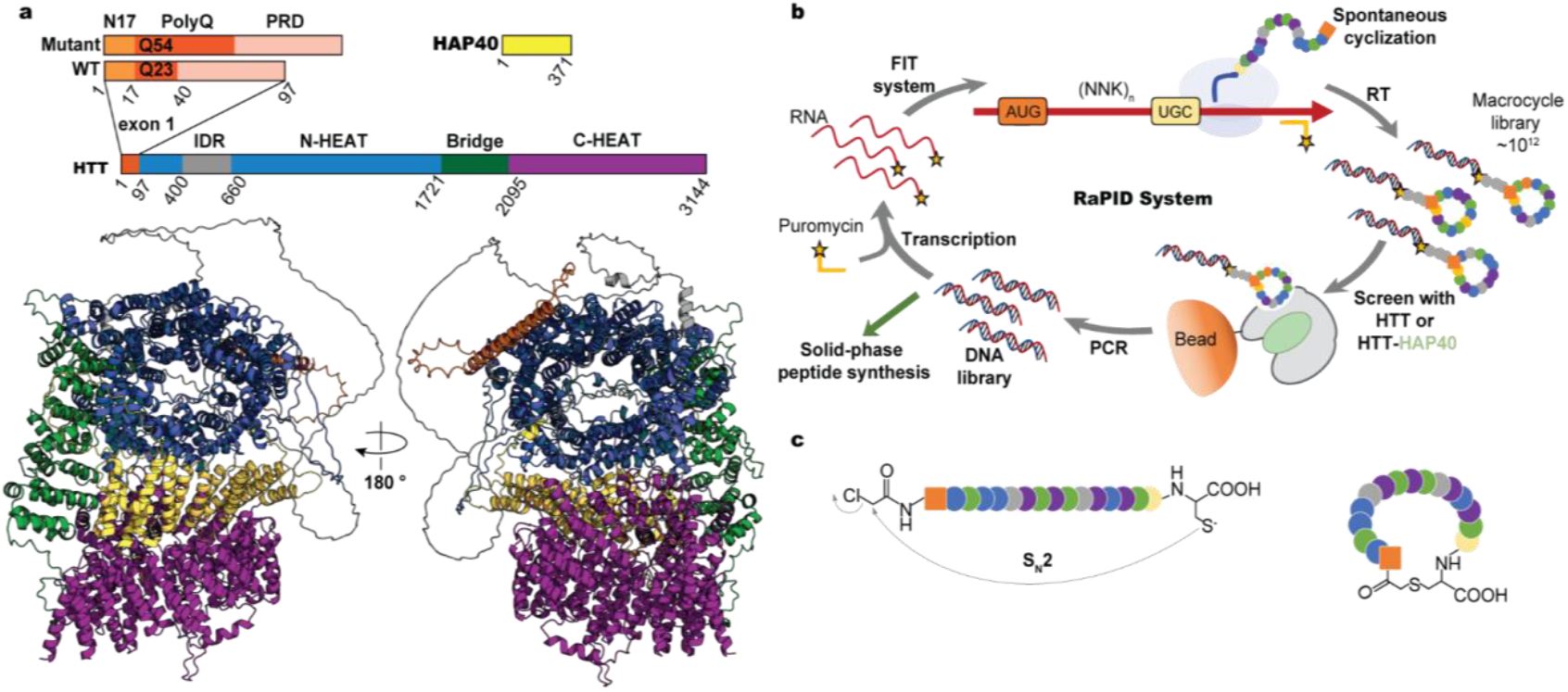
Macrocycle discovery and selection. **a**, Structure diagram of HTT-HAP40. HTT contains the N17, polyglutamine (polyQ), and proline rich domain (PRD), as well as the globular N-HEAT (blue), Bridge (green), and C-HEAT (purple) regions. A large intrinsically disordered region (IDR – grey) interrupts the N-HEAT domain. The wildtype (WT) protein in this study is Q23 (3144 aa), and the mutant is Q54. HTT forms a stable noncovalent heterodimer with HAP40 (pastel green). An AlphaFold3 structure of full-length HTTQ23-HAP40 was used here model the full-length proteins. **b**, RaPID is an RNA-display technique coupled to FIT, facilitating the incorporation of nonstandard residues which enable macrocyclization. **c**, Spontaneous cyclization of peptides enabled by nonstandard residue incorporation at the N-terminus, such as chloroacetyl-Tyr. Adapted from Saha, Suga, and Brik (2023) (Saha et al., 2023).

Immobilized HTTQ23 and HTTQ23-HAP40 were used to select binders from the theoretical ~10^12^ macrocycle pool. Following 5-6 rounds of screening, we observed enrichment for several sets of converging peptide sequences. To identify enantioselective ligands, we compared results L and D amino acid macrocycle libraries (Imanishi et al., 2021). Peptide sequences that selected from both D and L libraries were discarded. The remaining enriched enantioselective peptides were aligned by sequence and ranked based on enrichment (**Supplementary Figure 1**).

### Biophysical characterization of HTT-macrocycle interactions

The highest enriched and sequence converged macrocycles, named HD1-HD6, HL1-HL6, HHD1-HHD8, and HHL1-HHL7 (*where H = HTT and HH = HTT-HAP40 selected binders, and L and D indicate the initiating Tyr enantiomer*), had distinct sequences, suggestive of sampling a variety of epitopes on HTT and HTT-HAP40. These 27 macrocycles were synthesized by solid-phase peptide synthesis without tags for *in vitro* characterization. Five macrocycles, HL2 (YTARYLTLGTLHYKC), HL5 (YTHFQVAPWIQSLC), HD4 (^D^YDIWCETNKQTGILVLC), HHL1 (YLIRSSFNWFVFVEVPC), and HHD3 (^D^YTRHWQNNWWILYHEFC), had low nanomolar affinity for HTT or HTT-HAP40 upon initial screening by surface plasmon resonance (SPR) (**Figure 2b, Supplementary Figure 2**). HL5 did not engage HTTQ23-HAP40, while HL2 and HD4, despite RaPID selection using HTT, interacted with both HTT and HTTQ23-HAP40. This suggested HL2 and HD4 likely bound HTT on a surface available in both apo HTT and the heterodimer HTT-HAP40 complex.

**Figure 2.**
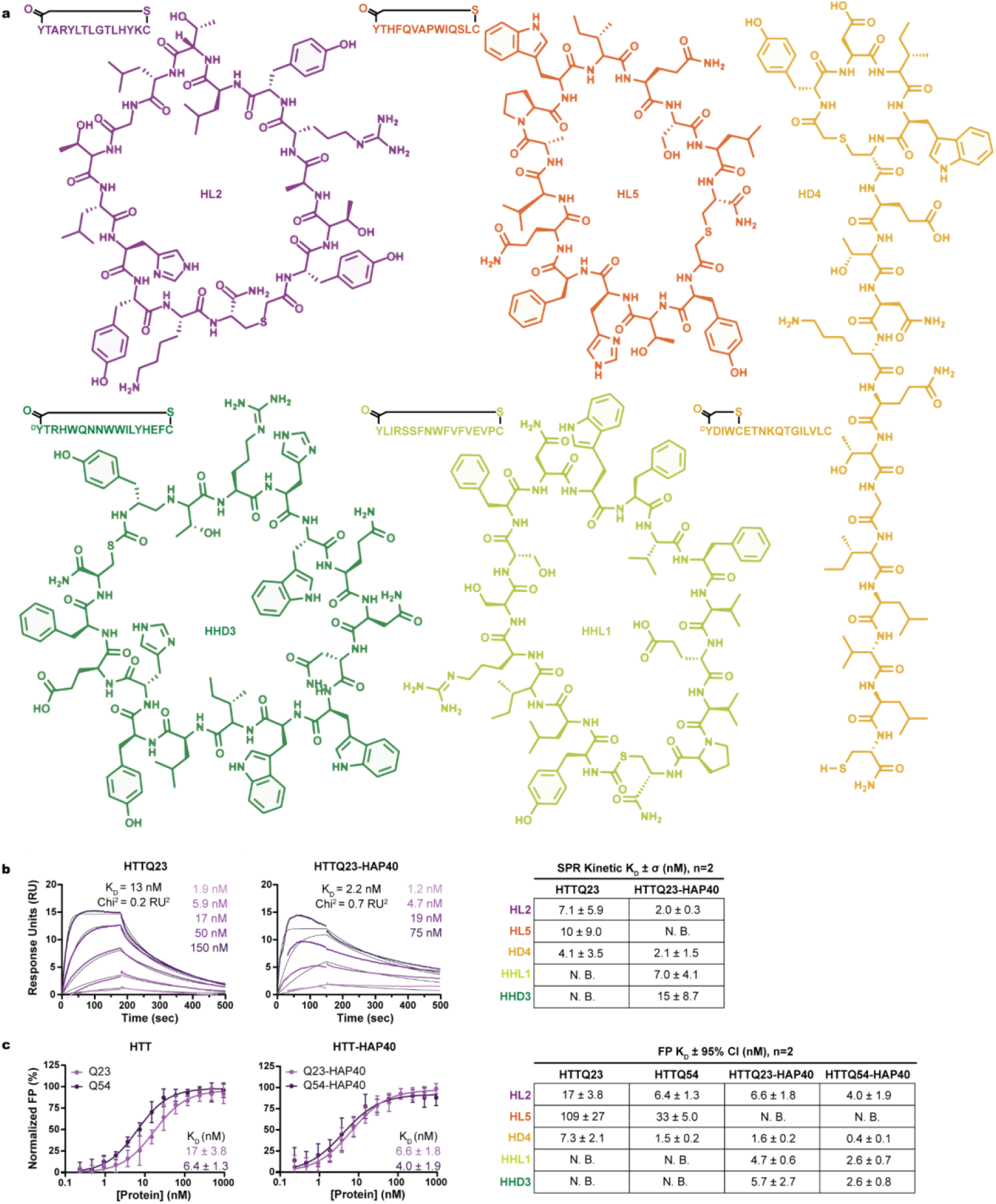
Macrocycle identity and affinity measurements. ***a***, *Sequence and structure of macrocycles validated by SPR. Assessment of HTTQ23 and HTTQ23-HAP40 binding by*, ***b***, *SPR and*, ***c***, *FP of exemplary macrocycle HL2 and summary tables for remaining macrocycles. Detailed affinity measurements for all macrocycles by SPR and FP available in* ***Supplementary Figure 2***. Both SPR and FP results indicate that HHL1 and HHD3 require the presence of HAP40 to interact with HTT. By contrast, HL5 interacts with apo HTT more tightly than HTT-HAP40 suggesting a potential binding site proximal to the interface between HTT and HAP40. Both HL2 and HD4 bound apo or HTT-HAP40, suggested a binding pocket distal to the HTT-HAP40 interface. Notably, significant selectivity for polyQ length was not observed since affinities fell into a similar range, suggesting the N-terminal polyQ tract did not play a major role in binding.

The peptide C-terminus of macrocycles discovered using RaPID can be derivatized with minimal risk of binding abrogation. We made use of this feature to generate C-terminally fluorescein isothiocyanate (FITC)-labeled macrocycles. These were used in a fluorescence polarization (FP) assay to quantify their affinity for solution-phase HTT or HTT-HAP40 with Q23 or Q54, representative of wildtype and disease-state proteins (**Figure 2c**). While the tightest binding indicated by dissociation constant (K_D_) was observed for HD4 (K_D_ = 0.4 nM with HTTQ54-HAP40), the weakest was observed for HL5 (K_D_ = 109 nM for HTTQ23).

We further verified the specificity of binding of the macrocycles for HTT and HTT-HAP40 by displacement assay (**Supplementary Figure 3**) with FITC-labeled and unlabeled macrocycle pairs. All labelled macrocycles were displaced by their sequence-matched unlabeled macrocycle, with IC50 values in the nanomolar range. HHD3 displaced fluorescently labeled HHL1, suggesting they bind at similar or overlapping sites.

### Mapping of macrocycle-HTT binding interfaces using FP

To further localize the binding of each macrocycle, we assessed their binding in an FP assay with HTT subdomains: NTD (aa. 97-2069), CTD (aa. 2095-3138), and CTD-HAP40 (**Figure 3a**) (Alteen et al., 2023). We previously demonstrated these domains are functional and can reconstitute the intact full-length complex with HAP40. Notably, our NTD construct lacks the polyQ tract and, unlike CTD, cannot be co-purified with HAP40 as a stable complex.

**Figure 3.**
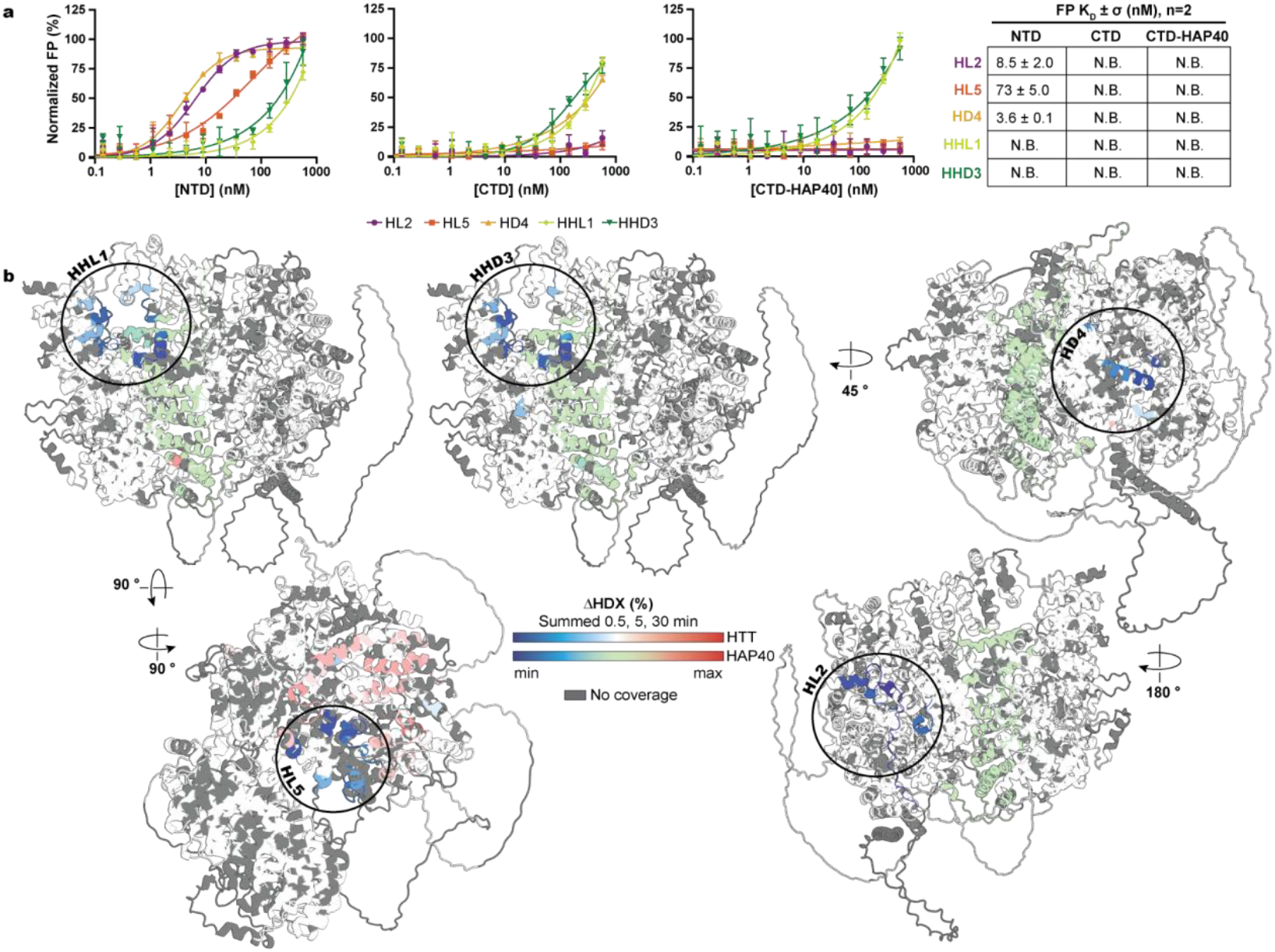
Mapping macrocycle binding interfaces. **a**, Determination of macrocycle affinity for HTT structure-rationalized subdomains by FP. Each plot represents a technical triplicate of each macrocycle tested against HTT’s CTD, NTD, and CTD-HAP40. The summary table reports the mean of a biological duplicate (n=2) ± σ. N.B. indicates no binding could be accurately estimated or detected at the concentrations tested. **b**, Differential Hydrogen-Deuterium Exchange Mass Spectrometry (ΔHDX-MS) of HTTQ23-HAP40 + HHD3, HHL1, HD4, or HL2, and HTTQ23 + HL5. Cumulative differences (0.5, 5, 30 mins) in fractional uptake exceeding cumulative error are shown on the HTT-HAP40 A3F model as increases (red), decreases (blue), statistically insignificant (white/green), and no coverage (grey).

Strong binding was observed between the NTD and HL2 (8.5 nM), HD4 (3.6 nM), and HL5 (73 nM). This binding occurred in the absence of HAP40, and with a similar binding constant to full-length HTT proteins, consistent with a binding pocket outside of the HTT-HAP40 heterodimerization interface. Notably, since the NTD construct lacks a polyQ tract, these data are indicative of polyQ independent binding, further supporting the lack of selectivity for Q23 or Q54. HHL1 and HHD3 binding affinity was weak or exceeded the concentrations of HTT subdomains tested. These data suggest that HHL1 and HHD3 may bind at or near the NTD-HAP40 interface, a surface not present in the subdomain constructs.

### Mapping of macrocycle-HTT binding interfaces using hydrogen-deuterium exchange mass spectrometry

Differential hydrogen-deuterium exchange mass spectrometry (ΔHDX-MS) (Wolf et al., 2024) was performed to more precisely map macrocycle binding interfaces with either apo HTT or the HTT-HAP40 complex (see **Supplementary Figures 4-8** for peptide coverage, bar plots, and the per-residue heat maps). The cumulative ΔHDX over 0.5, 5 and 30 minutes is illustrated in **Figure 3b** using the AlphaFold3 model for HTT-HAP40 (Abramson et al., 2024), highlighting the impact of macrocycle binding on HTT and HTT-HAP40 conformational dynamics.

Macrocycles HHD3 and HHL1 appeared to occupy the same binding site, formed by a ring of residues at the intersection between HTT-NTD and CTD, and HAP40: HTT aa. 1992-2005, 3004-3013, 3047-3060, 3074-3092 and HAP40 C-terminal aa. 261-303. Minimal localized changes were also observed in HAP40 N-terminal region, increasing (aa. 65-68) and decreasing (aa. 112-120) in deuterium uptake for HHL1 and HHD3, respectively, perhaps due to allosteric effects in HAP40. The HDX-identified binding sites of HHL1 and HHD3 were consistent with FP observations (**Figure 2bc** and **3a**), where a binding curve could not be generated in the absence of HAP40 and upon HTT truncation.

Consistent with HL2 and HD4’s nanomolar binding affinity for HTT, HTT-HAP40, and NTD constructs, both macrocycles induced decreased deuterium uptake at HTT’s NTD surface (**Figures 2 and 3**), in regions spanning aa. 387-408, 651-689, and 696-715 for HL2 and aa. 155-168, 1323-1326, 1363-1371, 1427-1443, and 1469-1473 for HD4. These regions would be accessible regardless of complexation with HAP40. Finally, HL5, seemingly destabilized HTT upon binding (**Figure 3**), causing broadly increased deuterium uptake across the NTD (aa. 791-799, 838-843, 903-907, 942-963, 1099-1111, 1126-1143, 1322-1326, 1360-1374, 1434-1443, 1474-1478, 1490-1500, 1512-1525, 1536-1552, and 1567-1577), and locally stabilized a ring of residues encompassing aa. 909-913, 963-967, 998-1004, 1015-1024, and 1034-1058, and 1071-1085 that would otherwise interface with HAP40. Consistently, no statistically significant cumulative HDX differences were observed when HL5 was incubated with HTTQ23-HAP40, suggesting this binding site was not readily accessible in the presence of HAP40 (**Supplementary Figure 9**). This was again consistent with HL5 binding apo HTT and the NTD subdomain construct in our FP assay. Thermal shift assays revealed that HL5 destabilized HTTQ54, while HHL1 and HHD3 stabilized HTTQ23-HAP40 in a dose-dependent manner, consistent with HDX-MS data (**Supplementary Figure 10**). No significant thermal effects were observed for HL2 or HD4, aligning with their peripheral binding sites on HTT.

### High resolution cryo-EM structure determination of HTT-macrocycle complexes

Cryo-EM was used to precisely determine the mode of macrocycle binding to HTT-HAP40. Informed by our crude mapping macrocycle binding sites, we determined the structures of HTT-HAP40 bound to macrocycles HHL1, HL2, and HD4 (Complex1), and of HTT-HAP40 bound to HHD3, HL2 and HD4 (Complex2). These were resolved to 2.1 Å and 2.3 Å respectively (see **Supplementary Table 1** for structure parameters, **Supplementary Figure 11** for data flow and **Supplementary Figure 12** for map-model fit). The structures revealed that macrocycles HL2 and HD4 bind exclusively to HTT, while HHL1 and HHD3 bind at the interface of the HTT-HAP40 complex, this is consistent with the SPR, FP, and HDX data. Each macrocycle engages the protein(s) through hydrogen bonds and salt bridges over a binding interface of ~ 750-1200 Å^2^ (**Figure 4, Supplementary Figure 13, Supplementary Table 2**).

**Figure 4.**
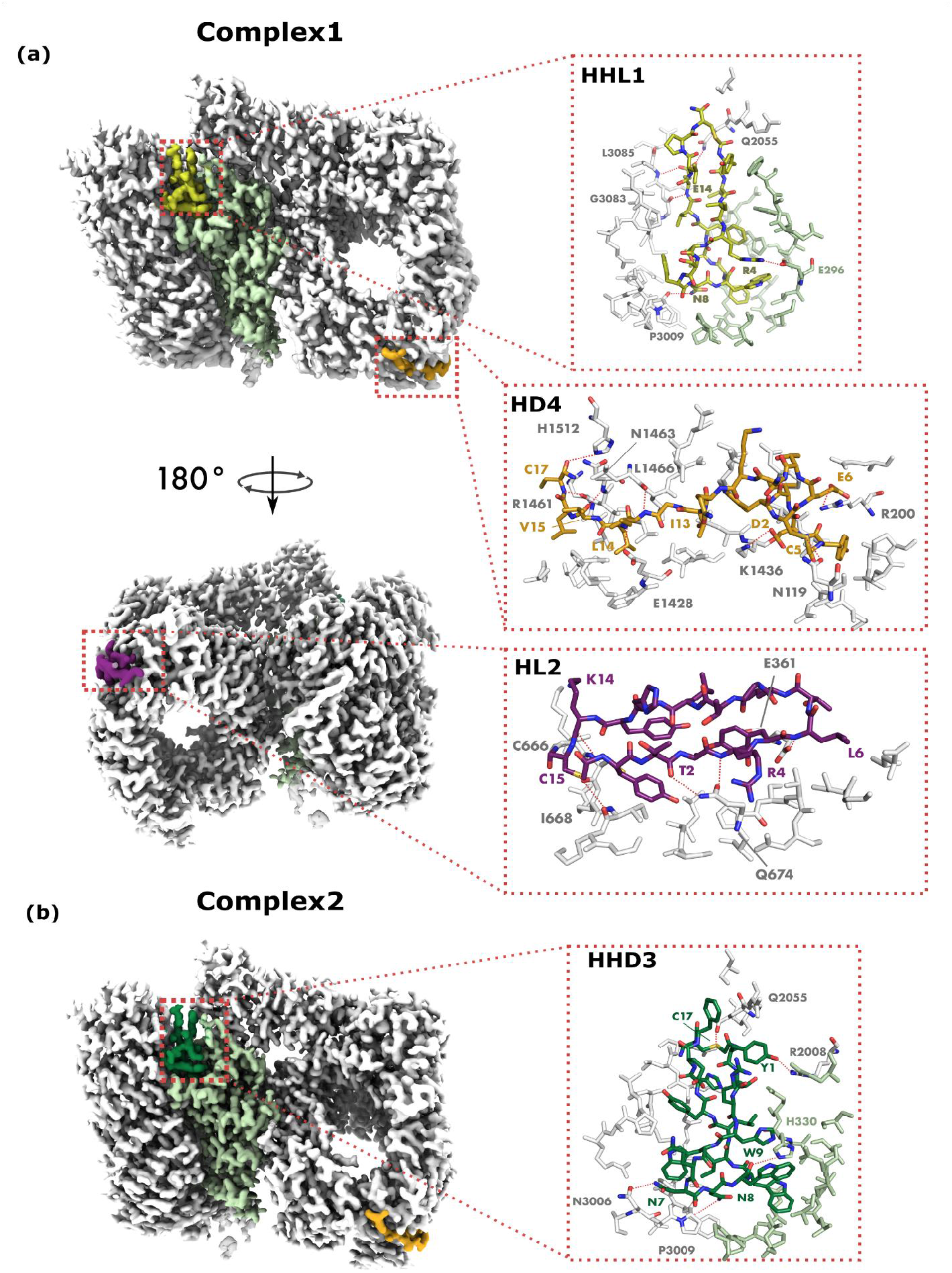
Cryo-EM resolved binding interactions of HL2, HD4, HHL1, and HHD3. **a**, Full complex of HTT-HAP40 and macrocycles HHL1, HD4 and HL2, showing binding interface and binding interactions as red dashed lines. **b**, HTT-HAP40 complex bound to HHL1, HD4 and HL2 showing interface and binding interactions.

#### HL2 interactions

HL2 binds a pocket in the HTT N-terminal domain. The pocket has strongly negative electrostatic potential, while HL2 is positively charged. The binding site is corroborated by the footprint observed by HDX-MS, and taken together, these data suggest that HTT residues 696-715 of an adjacent, predominantly solvent accessible loop, undergo reduction in orthosteric conformational dynamics. Notably, the macrocycle forms a beta-turn structure, where one arm of the turn is buried and the other is largely solvent exposed.

#### HHD3 and HHL1 interactions

As inferred from the HDX-MS data, HHD3 and HHL1 bind to the same pocket, which is at the HTT-HAP40 interface. This pocket is positively charged at the HTT interface and negatively charged at the HAP40 interface, and both HHL1 and HHD3 have a positively charged surface docking into the HAP40 interface. Interestingly, both HHL1 and HHD3 bind by bending into a ‘figure 8’ conformation.

#### HD4 interactions

HD4 binds a pocket that is also strongly negatively charged like the HL2-binding pocket, though the residues forming this pocket are primarily in the N-HEAT domain. This binding site is also in agreement with the HDX-MS data.

HL5 binds apo HTT but did not show significant binding to HTT-HAP40. Apo HTT proved to be an intractable target for high-resolution cryoEM, which has precluded the gathering of any atomic resolution structural information on the HTT-HL5 interaction.

### Macrocycles selectively bind endogenous HTT proteins from cells

Our structural and biophysical data demonstrated unequivocal binding to purified HTT. To explore binding and selectivity within a more physiological context, we used biotinylated derivatives of macrocycles to affinity purify HTT from cellular extracts, which we term “macrocycle precipitations” (MPs)(**Figure 5a**). The macrocycles were joined to the biotin using either a 2 amino acid linker (AK-Bio) or a PEG11 spacer (PEG11-Bio) (Supplementary Figure 12a). Extracts were prepared from HEK293T cells, including HTT knock-out lines and streptavidin beads were used to attempt to capture macrocycle–HTT complexes, and potentially associated binding partners.

**Figure 5.**
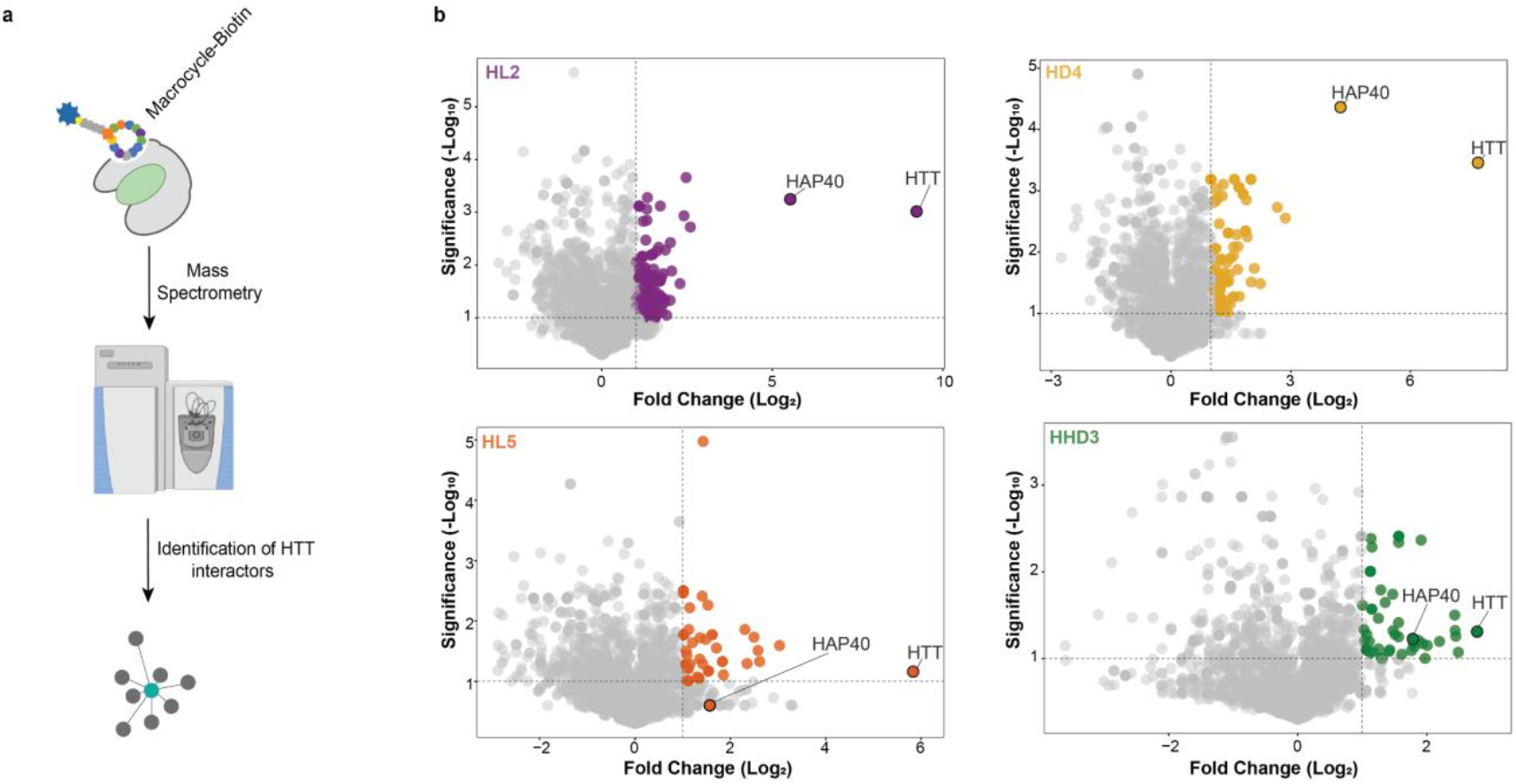
Macrocycles selectively bind endogenous HTT in MP assays. **a**. Schematic representation of the macrocycle precipitation (MP) assay workflow. HEK293T cells were lysed, and macrocycles were added to the lysate to bind to endogenous HTT in complex with other binding partners. Streptavidin beads were then introduced to capture macrocycle-bound complexes. After incubation, the beads were washed to remove unbound proteins, and HTT complexes were eluted for downstream analysis. **b**. Volcano plot of MP results showing log2 fold change (FC) and −log10 P values derived from normalized protein spectral counts

HTT, endogenously expressed in HEK293 cells, was shown to be captured by immobilized HD4-AK-Bio and HHD3-AK-Bio and by immobilized HL2-PEG11-Bio and HL5-PEG11-Bio (**Supplementary Figure 14b**). HHL1-AK-Bio or HHL1-PEG11-Bio showed no detectable binding of endogenous HTT. Notably, although HHL1 and HHD3 bind the same pocket, HHD3 consistently showed stronger performance in pulldown assays. One possible explanation is that HHL1 has a faster off-rate than HHD3, as indicated by the SPR sensorgrams (**Supplementary Figure 2b)**. It is possible that HTT dissociates from HHL1 during the dilution steps of the pulldown assay, resulting in reduced or short-lived target engagement. Alternatively, HHD3 may adopt a more favorable conformation within the binding pocket, resulting in improved target engagement.

To assess macrocycle selectivity and identify potential HTT interacting proteins, we performed LC-MS/MS analysis of the proteins in the MP eluates. To control for non-specific binding, we performed MPs using lysates from a HTT CRISPR knockout cell line (Jung et al., 2021). Across all macrocycles, HTT emerged as the most enriched protein after background subtraction (**Figure 5b**). Given we performed affinity purification using macrocycles targeting distinct HTT surfaces, we hypothesized that any real HTT protein binder might be co-purified with two or more macrocycles. HAP40 was consistently enriched in pulldowns using HL2, HD4, and HHD3, aligning with biophysical evidence that these macrocycles bind the HTT-HAP40 complex (**Supplementary Figure 14b**). In contrast, HAP40 was not enriched by HL5, consistent with its specificity for apo HTT.

### Robust Detection of the HTT-HAP40 Complex Suggests the Dominant Proteoform Across Contexts

Having demonstrated selective interactions between macrocycles and endogenous HTT and HAP40 proteins in HEK293T cells, we next sought to determine whether this interaction is preserved in models with greater relevance to HD biology. We extended our analysis to isogenic neuronal progenitor cells (NPCs), which more closely resemble the neuronal populations affected in HD, both functionally and in terms of gene expression profiles (Ooi et al., 2019). For these experiments, we selected the HL2 macrocycle based on its superior performance in HEK293T cells, where it exhibited the highest pulldown efficiency and strongest enrichment for HTT. Thus, we compared HL2 protein enrichment profiles across isogenic NPC lines harboring non-expanded (Q30/Q19) and expanded CAG repeats (Q45/Q19 and Q81/Q27).

Successful macrocycle-mediated pulldown was first confirmed by western blot in all NPC lines (**Supplementary Figure 14a**). Subsequent LC-MS/MS analysis revealed that HTT remained the most abundantly enriched protein across all lines, alongside HAP40, further supporting the conserved and interdependent nature of the HTT-HAP40 complex in a disease-relevant cell type (**Figure 6**). HTT and HAP40 were the only proteins consistently overlapping across all three NPC lines (**Supplementary Figure 14b**).

**Figure 6.**
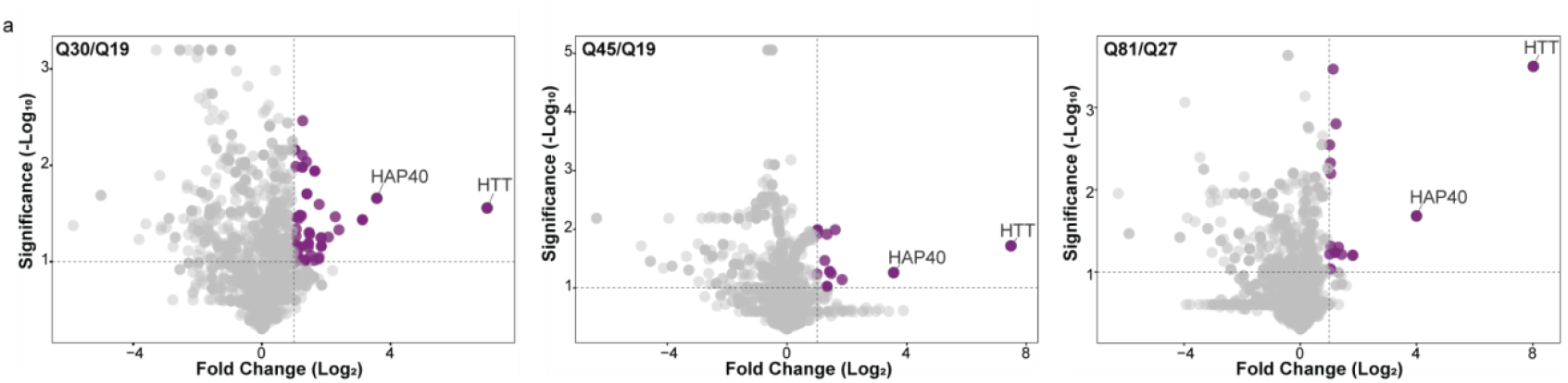
HTT and HAP40 form a constitutive complex across different polyQ lengths in NPCs. Volcano plot of HL2(PEG11) macrocycle pulldown experiments for Q30/Q19, Q45/Q19, and Q81/Q27 HTT-expressing NPCs. Results are shown in log_2_ fold change (FC) and −log10 p-values derived from normalized protein spectral counts.

Beyond observing HTT-HAP40 interactions across multiple macrocycles, with the exception of HL5, which has binding preference for apo HTT, HAP40 consistently emerged as the most prominent and high-confidence co-purifying protein. Even more compelling is the observation that across all pulldown experiments, the ratio of spectral counts between HTT and HAP40 remained approximately consistent. This likely indicates that HTT and HAP40 exist in a stable, stoichiometric complex within the cell.

## Discussion

Here we report a suite of well-characterized chemical tools that specifically bind full-length HTT in with both expanded and WT poly Q lengths, and in complex with HAP40. Our macrocycles, offering high affinity and selectivity for specific HTT proteo-states, constitute the first reported biophysically and structurally validated reagents to HTT and have the potential to considerably enhance our ability to dissect HTT function. The atomic-resolution structures of HL2, HD4, HHL1, and HHD3 lay the foundation for downstream mimetic development. For example, the ‘figure 8’ formed by HHL1 and HHD3 upon binding suggest two discernable subdomains that can be used to derive small molecule binders. The interaction sites of the macrocycles could also be a clue to the location of other potential protein-protein interactions. We examined endogenous HTT interactions using macrocycles that bind distinct regions of the full-length protein and using HTT knockout cell lines as negative controls—currently the most rigorous benchmark for pull-down experiments. Taken together, this level of characterization of the macrocycles as affinity reagents exceeds that of most HTT antibodies.

Our results also underscore the central role of the HTT-HAP40 complex in cells. A multitude of potential HTT interactors have been reported, with the total number exceeding 3,000 (Aaronson et al., 2021). We identified a total of 361 proteins, 311 of which were not previously reported in the HINT database (Patil & Nakamura, 2005). In our macrocycle pull down experiments, HAP40 was the only interactor consistently and significantly enriched with HTT across multiple cell lines. This observation aligns with previous reports showing that HAP40 levels track closely with HTT levels, suggesting a tightly regulated relationship (Harding et al., 2021b). These findings raise the possibility that HAP40 binding may be required for HTT to carry out its cellular functions. However, whether apo-HTT (unbound HTT) has an independent role remains unknown and warrants further investigation.

Beyond the use of our macrocycles as research affinity reagents, our tools, assays, and structural data will be highly enabling for the development of new HTT binders for imaging applications, including proximity-based spectroscopic methods and positron emission tomography (PET) tracers **(PMID: 38695673, 31967355)** (Dickmann et al., 2024; Fazio et al., 2020). Insert here about optimizing cell permeability and small molecule design. Finally, these macrocycles also present an opportunity to advance HTT-targeting degradation strategies (Li et al., 2019; Yamashita et al., 2020). They can serve as HTT-specific ‘warheads’ in PROTAC design. Notably, a recent study (PMID: 37876498) demonstrated a fully peptidic PROTAC by linking a macrocycle to an E3 ligase-binding C-degron. A similar strategy for future studies of this nature are underway.

## Materials and Methods

### Protein Expression and Purification

As described previously, (Alteen et al., 2023; Harding et al., 2019), HTT (Q23 or Q54), HTT-HAP40, and structure-rationalized HTT domains (C-HEAT res. V2095-V3138, NTD res. T97-M2069) underwent baculovirus-mediated expression in Sf9 insect cells. Purified protein was obtained using FLAG affinity chromatography and subsequent size-exclusion chromatography (SEC), followed by intact mass spectrometry (MS) mass validation. C-terminally biotinylated HTT (apo and in complex with HAP40) used in SPR and RaPID was generated by cloning in a C-terminal Avi-tag and followed by co-expression with BirA and supplementation with exogenous biotin.

### Screen for macrocyclic peptides that bind to HTT and HTT-HAP40

We employed the RaPID system to screen libraries of over 10^12^ unique macrocyclic peptides to identify candidate binders based on their ability to bind tightly to the mHTT or mHTT-HAP40. A puromycin-ligated mRNA library was constructed to encode peptides with N-chloroacetyl-L-Tyrosine (LY-library) or N-chloroacetyl-D-Tyrosine (DY-library) as the initiator amino acid, followed by a random peptide region consisting of 6-15 residues, a cysteine and ending with a short linker peptide. Upon translation of these mRNAs, the chloroacetyl group on N-terminus of the linear peptides spontaneously cyclizes with the downstream cysteine to form thioether-macrocyclic peptides. The cyclic scaffold ensured that the LY-library and the DY-library diversified three-dimensional structures. Each cyclic peptide was covalently linked to its corresponding mRNA template via the puromycin linker for later amplification and DNA sequencing. The peptide-oligonucleotides (mRNA/cDNA) fusions were incubated with streptavidin-conjugated magnetic beads to which a N-terminally biotinylated HTT or HTT-HAP40 and pulldown for 30 min at room temperature (negative selection against beads and wild type proteins). The unbound fraction was then applied to magnetic beads with mHTT or mHTT-HAP40 for selections. Following four or five rounds of selection, the recovery rate of peptide–mRNA fusion molecules was significantly increased, suggesting that the population of cyclic peptides binding HTT or HTT-HAP40 were selectively enriched. Sequence analysis of the respective enriched libraries yielded unique sequences for the peptides.

### Chemical synthesis of peptides

Macrocyclic peptides were synthesized by standard Fmoc solid-phase peptide synthesis (SPPS) using a Syro Wave automated peptide synthesizer (Biotage). The resulting peptide–resin (25 μmol scale) was treated with a solution of 92.5% trifluoroacetic acid (TFA), 2.5% water, 2.5% triisopropylsilane and 2.5% ethanedithiol, to yield the free linear N-ClAc-peptide. Following diethyl ether precipitation, the pellet was dissolved in 10 ml triethylamine containing DMSO and incubated for 1 h at 25 °C, to yield the corresponding macrocycle. The peptide suspensions were then acidified by addition of TFA to quench the macrocyclization reaction. The macrocycle was purified by RP-HPLC, using a Prominence HPLC system (Shimadzu) under linear gradient conditions. Mobile phase A (comprising water with 0.1% TFA) was mixed with mobile phase B (0.1% TFA in acetonitrile). Purified peptides were lyophilized in vacuo and molecular mass was confirmed by MALDI MS, using an AutoFlex II instrument (Bruker Daltonics).

### SPR Assay

0.4 mg/mL HTTQ23 and HTTQ23-HAP40 were immobilized on a Streptavidin or Biotin CAPture chip in SPR Running Buffer (10 mM HEPES pH 7.4, 150 mM NaCl, 1 mM EDTA, 0.005% Tween-20 (v/v), 2% DMSO (v/v)) to a response of 2000 and 2500 RU, respectively. Multi Cycle Kinetics were performed for HL2, HD4, HL5, HHD3, and HHL1. A 1:1 binding model was used to calculate kinetic dissociation coefficient (K_D_) and the fit of the theoretical curve was determined by Chi^2^ (RU^2^).

### FP Assay

In a low volume, flat bottom black polystyrene 384 well plate (Corning, REF 3820), 20 or 2000 nM HTTQ23 or HTTQ23-HAP40 were serially diluted (12-point, 2-fold) into assay buffer (20 mM HEPES pH 7.4, 150 mM NaCl, 1 mM TCEP, 0.005% Tween-20 (v/v), 0.2% DMSO (v/v)) in triplicate. 1 uL of 20 nM of C-terminally FITC labeled macrocycle (100% DMSO) was added to 19 uL of serially diluted protein. Immediately, the plate was covered with an aluminum foil seal and centrifuged at 4 °C, 1000 x g for 2 min. Fluorescence polarization (mP) was measured using a Synergy Neo2 Multi-Mode Assay Microplate Reader (SickKids Hospital, Structural and Biophysical Core Facility, Toronto, Canada) with excitation 485 nm, emission 528 nm, gain 100, and Top optics. The data were visualized using GraphPad Prism after transformation relative to the baseline macrocycle FP (0 nM protein) and normalization relative to the mean highest FP detected. The FP was fit using Specific binding with Hill slope.

### HDX-MS

15 μM HTTQ23 or HTTQ23-HAP40 (20 mM HEPES pH 7.0, 300 mM NaCl, 2.5% glycerol (v/v), 1 mM TCEP, 2% DMSO (v/v)) were prepared alone or with 1:2 macrocycle (or 1:4 in the case of HL5 to accommodate its weaker K_D_), and stored at 0 °C. Samples were labeled at 3 timepoints (0.5, 5, 30 mins) with 87% (v/v) deuteration buffer (10 mM Phosphate Buffer pD 7.5, 150 mM NaCl) at 20 °C. The HDX reaction was quenched at 50% (v/v) with 7.5 M Guanidine-HCl, 0.5 M TCEP, 100 mM Phosphate Buffer pH 2.5 at 0 °C for 2 min and subsequently diluted with an equal volume of quench dilution buffer (100 mM Phosphate Buffer, pH 2.5). The protein was digested using 1:1 Nepenthesin(II)-Pepsin (Affipro) followed by desalting (ACQUITY UPLC BEH C18 VanGuard Pre-column, Waters) and reverse-phase separation (ACQUITY UPLC BEH C18 Column, Waters). Liquid handling was achieved using the ACQUITY UPLC M-Class System with HDX Technology (Waters) and PAL3 Robotic Tool Change (LEAP, Trajan Scientific and Medical). The peptides were then analyzed using the Select Series Cyclic IMS mass spectrometer (Q-IMS-TOF, Waters). Peptide identification and HDX data analysis were achieved using ProteinLynx Global Server 3.0.3 (Waters) and DynamX 3.0 (Waters), respectively. Microsoft Excel and PyMOL 2.5 were used for visual presentation of HDX data. An AlphaFold 3 model was used to incorporate missing loops for depicting HTTQ23 and HTTQ23-HAP40. The peptide sequence coverage for HTT and HAP40 was 56.6-76.8% and 77.4-82%, respectively (**Supplementary Figure 3-5**). For statistical significance, the cumulative ΔHDX-MS signals exceeded the cumulative propagated error at triple the standard deviation (**Supplementary Figure 6-8**).

### Thermal Shift Assay

In an Axygen® 384-well PCR microplate (REF# PCR-384-LC480-W-NF, Corning), samples were mixed to yield a final concentration of 0.25 mg/mL Q54 or Q23/HAP40, a 5-pt 2-fold serial dilution of each macrocycle (top concentration of 10 µM, 2.5% DMSO), and 5X SYPRO™ Orange (CAT# S6650, Thermo Fisher Scientific) in buffer (20 mM HEPES pH 7.4, 150 mM NaCl, 1 mM TCEP, 0.005% Tween-20). Differential Scanning Fluorimetry (DSF) was performed using a LightCycler® 480 (Roche LifeSciences) set to a 4 °C/min temperature ramp (20-95 °C) and fluorescence detection at 580 nm. Melting temperatures (T_m_) were obtained from the minima of the negative derivative of each curve. The propagated error of the thermal shift (ΔT_m_) was calculated from the standard deviation of each T_m_ mean (n=3). To ensure a maximally macrocycle-bound fraction (K_D_ ~10 nM; ~100% bound) at 0.25 mg/mL HTT or HTT-HAP40 (~0.7 µM), the upper limit of macrocycle tested was 10 µM.

### Cryo-EM sample preparation and data acquisition

HTT-HAP40 macrocycle complexes were diluted to 0.2 mg/ml in 25 mM HEPES pH 7.4, 150 mM NaCl, 0.025 % w/v CHAPS, and adsorbed onto lightly glow-discharged (10 s, 15 mA) suspended monolayer graphene grids (Graphenea) for 60 s. Grids were then blotted with filter paper for 1 s at 100 % humidity, 4 °C and frozen in liquid ethane using a Vitrobot Mark IV (Thermo Fisher Scientific). Movies were collected in counted mode, in Electron Event Representation (EER) format, on a CFEG-equipped Titan Krios G4 (Thermo Fisher Scientific) operating at 300 kV with a Selectris X imaging filter (Thermo Fisher Scientific) and slit width of 10 eV, at 165,000x magnification on a Falcon 4i direct detection camera (Thermo Fisher Scientific), corresponding to a calibrated pixel size of 0.732 Å. Movies were collected at a total dose of 51.8 e-/Å^2^ (Table 1), fractionated to ~ 1.0 e^−^/Å^2^ per fraction for motion correction.

### Cryo-EM data processing and modeling

Patched motion correction, CTF parameter estimation, particle picking, extraction, and initial 2D classification were performed in SIMPLE 3.0(Caesar et al., 2020). All downstream processing was carried out in cryoSPARC (Punjani et al., 2020) or RELION 3.1 (Zivanov et al., 2019),using the csparc2star.py script within UCSF pyem(Asarnow et al., 2019) to convert between formats. Global resolution was estimated from gold-standard Fourier shell correlations (FSCs) using the 0.143 criterion and local resolution estimation was calculated within cryoSPARC. The cryo-EM processing workflow for the HTT-HAP40-HHL1-HL2-HD4 complex is outlined in **Supplementary Figure 10a-c**. Briefly, downsampled particles (1.464 Å / pixel, 178 × 178 pixel box size) were subjected to two rounds of reference-free 2D classification (k=200 each) within cryoSPARC. Selected particles (544,842) were subjected to multi-class *ab initio* reconstruction using a maximum resolution cutoff of 6 Å, generating four volumes. Particles (230,039) from the most populated and structured class were selected and non-uniform refined against their corresponding volume lowpass-filtered to 8 Å, generating a Nyquist-limited 3.0 Å map. Bayesian polishing was then performed in RELION back to the nominal movie pixel size of 0.732 Å followed by extraction in a box size of 512 × 512 pixels. Polished particles were non-uniform refined against their corresponding upsampled volume while fitting per-particle CTF parameters and global beam tilt and trefoil, resulting in a 2.1 Å volume. This volume was sharpened by the highres model of deepEMhancer (Sanchez-Garcia et al., 2021). The cryo-EM processing workflow for the HTT-HAP40-HHD3-HL2-HD4 complex is outlined in **Supplementary Figure 10d-f**. Briefly, downsampled particles (1.464 Å / pixel, 178 × 178 pixel box size) were subjected to two rounds of reference-free 2D classification (k=200 each) within cryoSPARC. Selected particles (715,518) were subjected to multi-class *ab initio* reconstruction using a maximum resolution cutoff of 6 Å, generating four volumes. Particles (211,814) from the most populated and structured class were selected and non-uniform refined against their corresponding volume lowpass-filtered to 8 Å, generating a Nyquist-limited 3.0 Å map. Bayesian polishing was then performed in RELION back to the nominal movie pixel size of 0.732 Å followed by extraction in a box size of 512 × 512 pixels. Polished particles were non-uniform refined against their corresponding upsampled volume while fitting per-particle CTF parameters and global beam tilt and trefoil, resulting in a 2.3 Å volume. This volume was sharpened by deepEMhancer (Sanchez-Garcia et al., 2021) to obtain the final high-resolution map.

The previously determined structure of the HTT-HAP40 complex (PDB 6×90) was used as an initial model for model building, and the macrocycles were built in *ab initio*. Cyclization of the macrocycles and energy minimization was done using Chimera (Pettersen et al., 2021). This initial model was then further refined using both the refinement and validation tools available in Coot (Emsley et al., 2010), and ISOLDE (Croll, 2018) within ChimeraX (Pettersen et al., 2021). The model was validated using Coot and MolProbity (Chen et al., 2010). Figures were generated using Chimera (Pettersen et al., 2004), ChimeraX and Pymol (Schrödinger, LLC, 2015).

#### Macrocycle Bead Conjugation

Macrocycles were pre-conjugated to streptavidin beads (Cytiva, 17-5113-01) by incubating with a 30 μL slurry of beads per macrocycle purification (MP) in Lysis Buffer (10 mM Tris-HCl, pH 7.9, 150 mM NaCl, 0.1% NP-40, 1:500 protease inhibitor cocktail (Sigma-Aldrich), 1:1,000 benzonase nuclease, and 2 mM MgCl_2_) for 1 hour at 4°C. A concentration of 1.25 μM macrocycle was used for conjugation. After incubation, beads were washed three times with 500 μL Lysis Buffer to remove unbound macrocycles and resuspended in 500 μL of lysate at 4 mg/mL protein concentration.

#### Cell Lysis

Cells were harvested by centrifugation at 1,500 rpm for 5 minutes, washed twice with PBS, and snap-frozen. Cell pellets were then resuspended in 500 μL of Lysis Buffer (as described above) and incubated on an end-over-end rotator at 4°C for 1 hour. Lysates were clarified by centrifugation at 20,000g for 15 minutes at 4°C, and the supernatant was transferred to fresh 1.5 mL Eppendorf tubes. Protein concentration was determined by BCA assay (Thermo Fisher Scientific, 23225), yielding approximately 4 mg of protein on average. Lysates were diluted to a final concentration of 4 mg/mL for use, providing 2 mg of total protein per MP.

#### Macrocycle Precipitation (MP) of HTT

Macrocycle-conjugated beads were incubated with 2 mg of lysate in Lysis Buffer for 1 hour at 4°C with rotation. After incubation, beads were collected by centrifugation at 3,000 rpm, and the flow-through was saved for analysis. Beads were then washed sequentially with 1 mL of Wash Buffer 1 (10 mM Tris-HCl, pH 7.9, 100 mM NaCl, 0.1% NP-40) and twice with Wash Buffer 2 (Wash Buffer 1 without detergent). Lastly, bound proteins were eluted from the beads with 200 μL of 0.5 M NH_4_OH and flash-frozen in liquid nitrogen.

#### Preparation and Analyses of MP Samples for Liquid Chromatography-Tandem Mass Spectrometry (LC-MS/MS)

All sample preparation, LC MS/MS data collection, and MS data searches were carried out at the SPARC BioCentre at The Hospital for Sick Children (Toronto, ON, Canada). Samples were reduced with DTT (10 mM, 60°C, 1 hour), alkylated with iodoacetamide (20 mM, room temperature, 45 min, dark), and digested overnight at 37°C with trypsin (2 mg; Pierce). Peptides were dried by vacuum centrifugation, desalted on C18 ziptips (Millipore) using a DigestPro MSi (Intavis Bioanalytical Instruments), and dried again by vacuum centrifugation before resuspension in Buffer A (0.1% formic acid (v/v) in water). Samples were analyzed by liquid chromatography tandem mass spectrometry (LC-MS/MS) using an EASY-nanoLC 1200 system with a 1 h analysis and an Orbitrap Fusion™ Lumos™ Tribrid™ Mass Spectrometer (Thermo Fisher Scientific). The LC portion of the analysis consisted of a 18 min linear gradient running 3-20% of Buffer A to Buffer B (0.1% formic acid (v/v) in acetone), followed by a 31 min linear gradient running 20-35% of Buffer A to Buffer B, a 2 min ramp to 100% Buffer B and 9 min hold at 100% Buffer B, all at a flow rate of 250 nL/min. Samples were loaded into a 75 mm x 2 cm Acclaim PepMap 100 Pre-column followed by a 75 µm x 50 cm PepMax RSLC EASY-Spray analytical column filled with 2 µM C18 beads (Thermo Fisher Scientific). MS1 acquisition resolution was set to 120000 with an automatic gain control (AGC) target value of 4 × 105 and maximum ion injection time (IT) of 50 ms for a scan range of m/z 375-1500, with dynamic exclusion set to 10 s. Isolation for MS2 scans was performed in the quadrupole with an isolation window of m/z 0.7. MS2 scans were performed in the ion trap with maximum ion IT of 10 ms, AGC target value of 1 × 104, and higher-energy collisional dissociation (HCD) activation with an NCE of 30.

MS raw files were analyzed using PEAKS Studio software (Bioinformatics Solutions Inc.) and Proteome Discoverer (version 3.1.1.93) and fragment lists searched against the human UniProt Reference database (uniprotkb_Human_All_UP000005640_20240726). For both search algorithms, the parent and fragment mass tolerances were set to 50 ppm and 0.02 Da, respectively, and only complete tryptic peptides with a maximum of three missed cleavages were accepted.

## Acknowledgments

We thank M. E. MacDonald and I. S. Seong from Massachusetts General Hospital for a gift of HTT null HEK293T cells. We thank Craig Simpson at SPARC BioCentre, Hospital for Sick Children, Toronto, Canada for his assistance with mass spectrometry mass spectrometry and associated data analysis.

