## Supplementary Information for "High-Affinity, Structure-Validated and Selective Macrocyclic Peptide Tools for Chemical Biology Studies of Huntingtin"

**Abstract**

Huntington's disease (HD) is a fatal neurodegenerative disorder caused by a CAG repeat expansion in the *Huntingtin* (*HTT*) gene, with no disease-modifying therapies currently available. The precise molecular function of the HTT protein is unclear, and the lack of selective chemical tools has limited functional studies. We have identified and characterized macrocyclic peptide binders targeting HTT. These binders exhibit low-nanomolar affinity *in vitro* and engage distinct HTT and HTT-HAP40 interfaces, as revealed by hydrogen-deuterium exchange mass spectrometry and cryo-electron microscopy. Chemoproteomics confirmed selective binding in cell extracts from wildtype but not HTT-null cell lines. HAP40 consistently and stoichiometrically co-purified with HTT across cell lines, including with HTT variants containing different CAG repeat lengths, highlighting the broad presence of the HTT-HAP40 complex.

### Supplementary Information

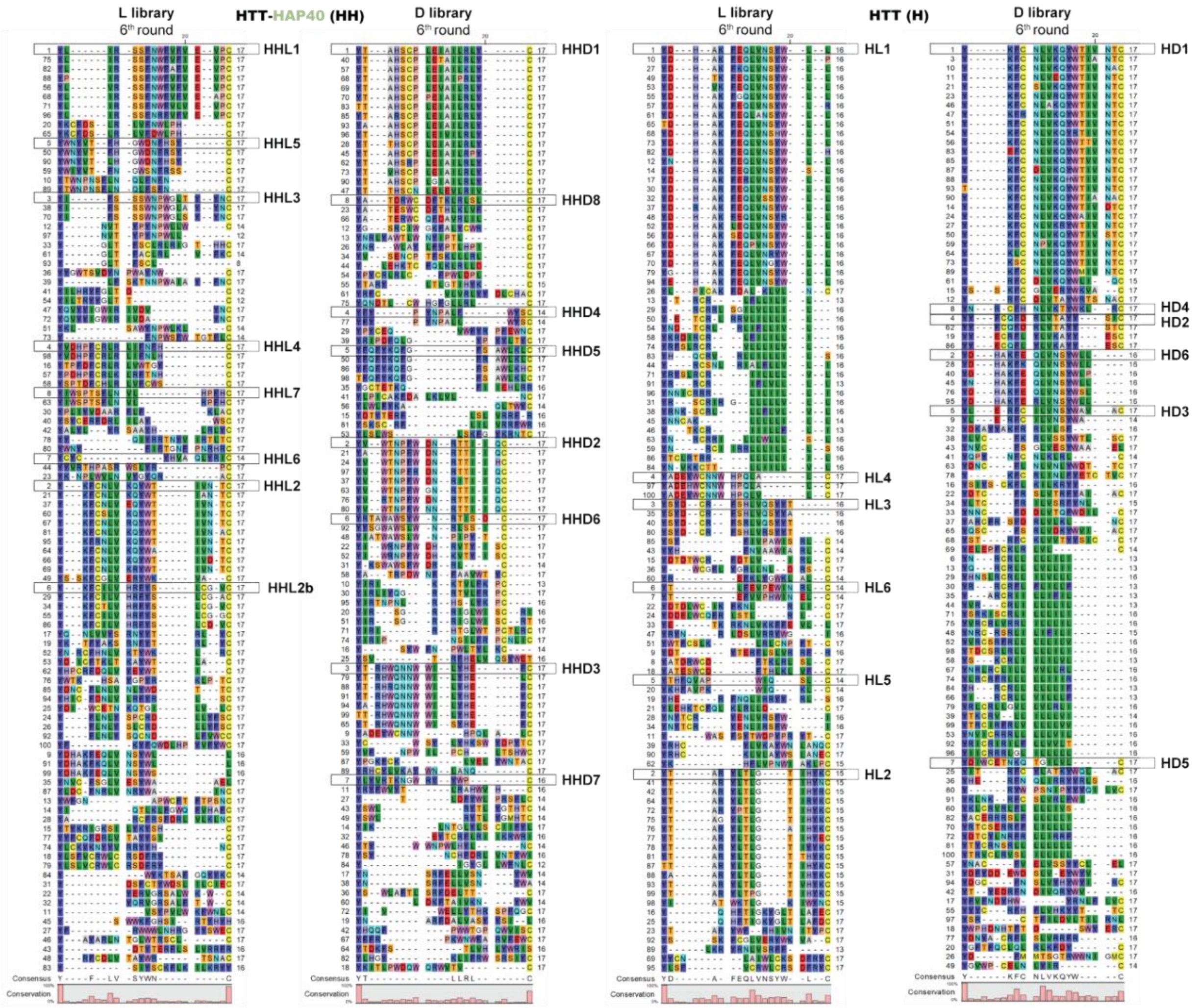


**Supplementary Figure 1.** Top 100 Aligned Peptide Sequences. Numbers to the left of each set indicate peptide enrichment rank (lower number corresponds to greater enrichment) while value on the right is the sequence length. Sequences aligned using ClustalX (PMID 18792934).

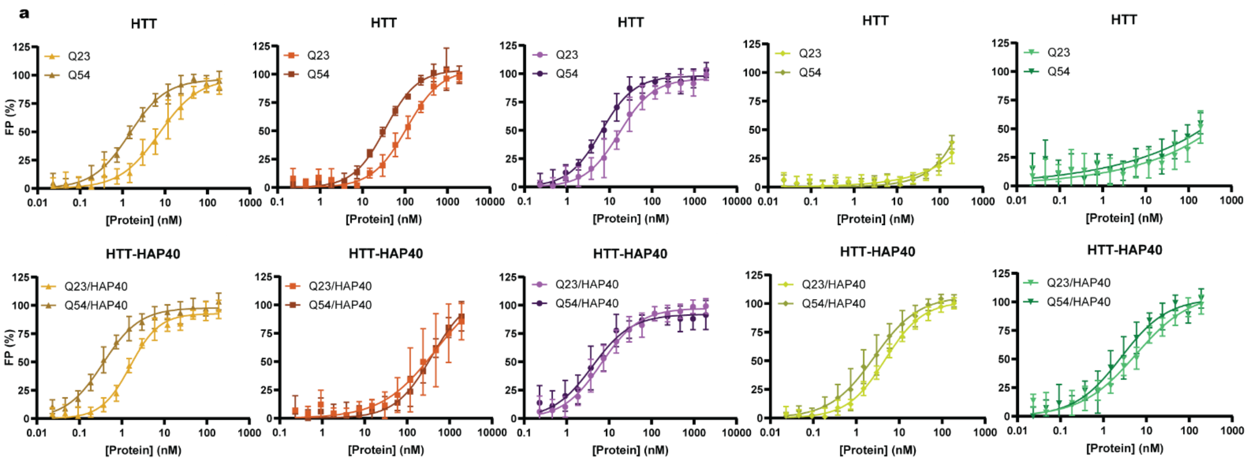


| | FP $K_D$ ± 95% CI (nM), n=2 | | | |
|---|---|---|---|---|
| | HTTQ23 | HTTQ54 | HTTQ23-HAP40 | HTTQ54-HAP40 |
| HL2 | 17 ± 3.8 | 6.4 ± 1.3 | 6.6 ± 1.8 | 4.0 ± 1.9 |
| HL5 | 109 ± 27 | 33 ± 5.0 | 462 ± undef. | 389 ± 676 |
| HD4 | 7.3 ± 2.1 | 1.5 ± 0.2 | 1.6 ± 0.2 | 0.4 ± 0.1 |
| HHL1 | No binding | No binding | 4.7 ± 0.6 | 2.6 ± 0.7 |
| HHD3 | No binding | No binding | 5.7 ± 2.7 | 2.6 ± 0.8 |

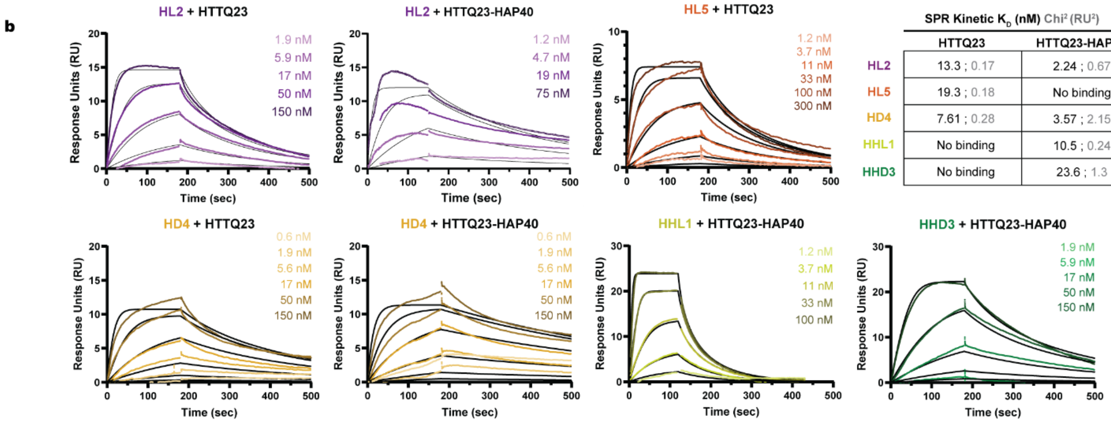


| | SPR Kinetic $K_D$ (nM) $Chi^2$ ($RU^2$) | |
|---|---|---|
| | HTTQ23 | HTTQ23-HAP40 |
| HL2 | 13.3 ; 0.17 | 2.24 ; 0.67 |
| HL5 | 19.3 ; 0.18 | No binding |
| HD4 | 7.61 ; 0.28 | 3.57 ; 2.15 |
| HHL1 | No binding | 10.5 ; 0.24 |
| HHD3 | No binding | 23.6 ; 1.3 |

**Supplementary Figure 2**. Fluorescence polarization assay and Surface plasmon resonance-derived affinity values for macrocycles and full-length HTT or HTT-HAP40. **a**, The FP-derived $K_D$ error was determined from the upper and lower limit of the 95% confidence interval of a biological duplicate (n=2). The FP dissociation coefficient ($K_D$) was determined using Specific Binding with Hill Slope. **b**, The SPR kinetic $K_D$ were determined by theoretical curve fit to the association (contact) and dissociation phases in the sensorgrams, and the $Chi^2$ represents the extent of variation between the theoretic and experimental curves.

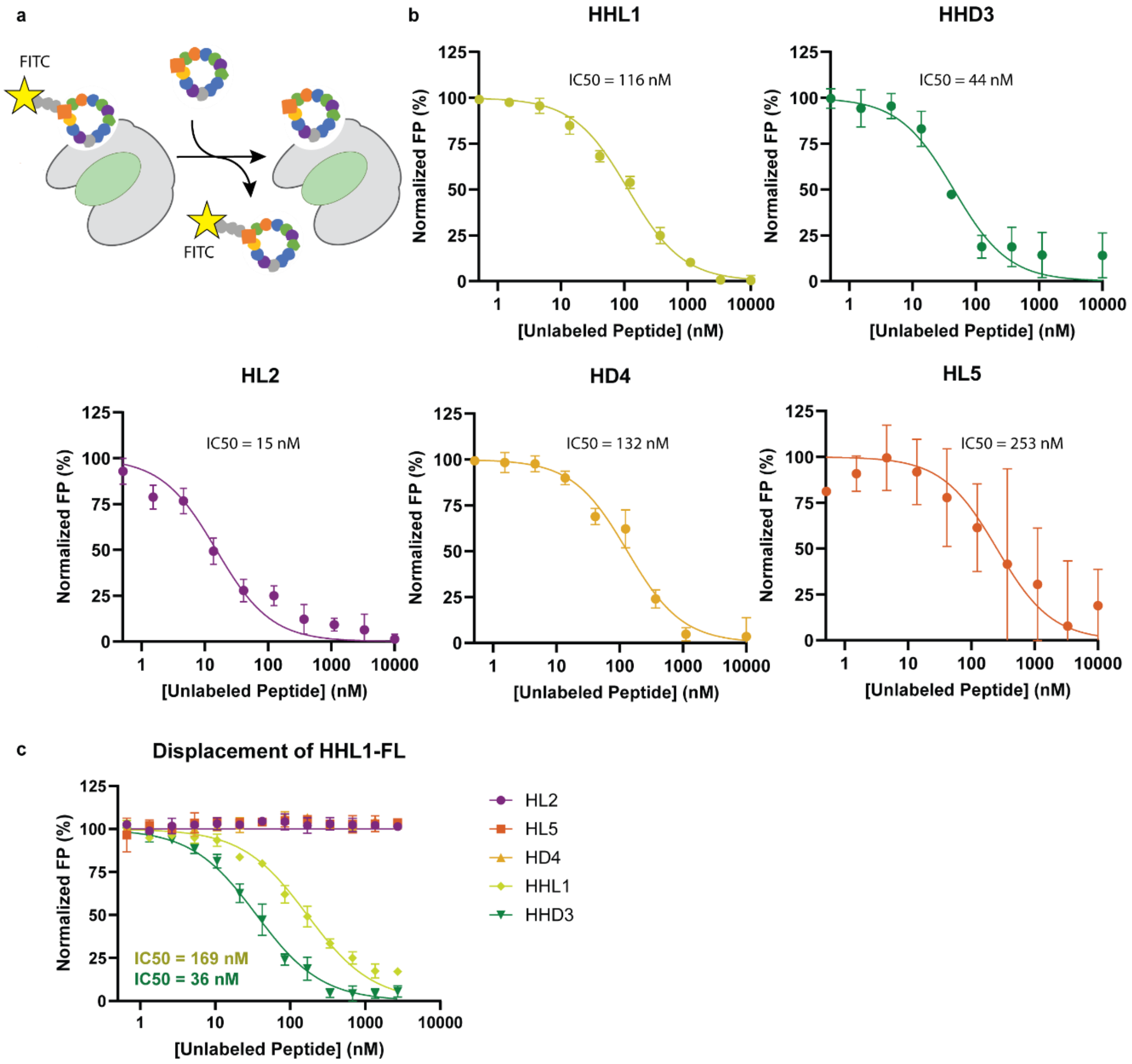


**Supplementary Figure 3.** Displacement of fluorescently labeled macrocycles. **a**, Schematic for assay. **b**, Each fluorescently labeled macrocycle is displaced by its unlabeled partner from HTT-HAP40 or HTT (HL5) Conducted as a biological singlet in technical triplicate. **c**, Fluorescently labeled HHL1 is displaced by itself and HHD3, but no other macrocycle.

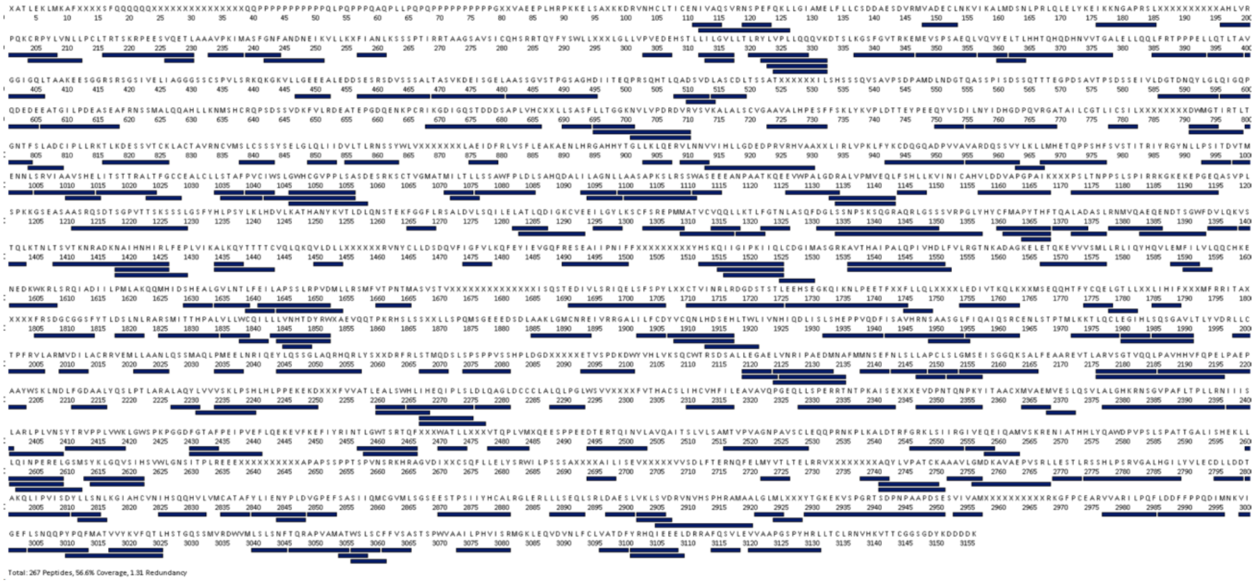


**Supplementary Figure 4.** Peptide Sequence Coverage of HTTQ23 for HL5 HDX-MS. 267 peptides, represented by navy blue rectangles, were analyzed to generate a sequence coverage of 56.6 % and a peptide-per-reside redundancy of 1.31.

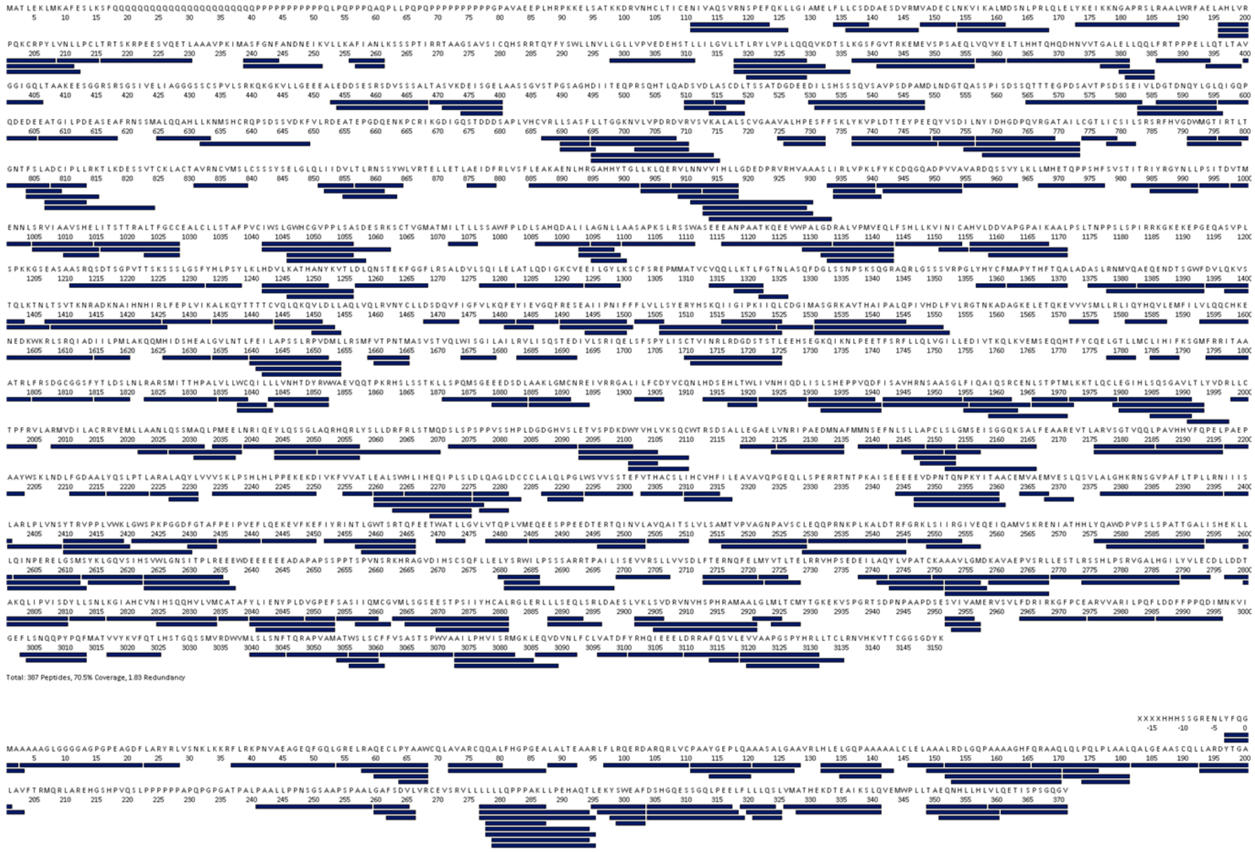


**Supplementary Figure 5.** Peptide Sequence Coverage of HTTQ23 and HAP40 for HHL1 and HHD3 HDX-MS. For HTTQ23 (HAP40), 387 (61) peptides, represented by navy blue rectangles, were analyzed to generate a sequence coverage of 70.5 % (77.4 %) and a peptide-per-reside redundancy of 1.83 (2.33).

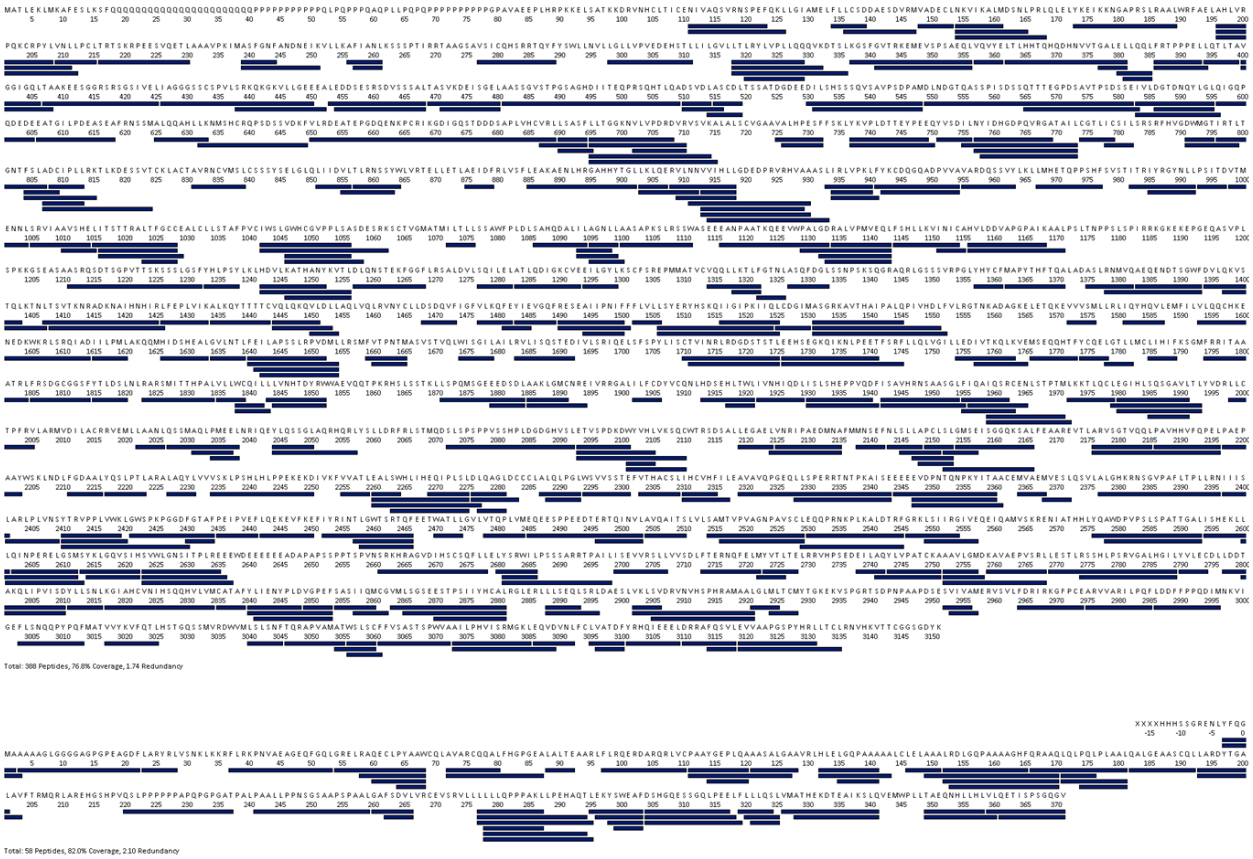


**Supplementary Figure 6.** Peptide Sequence Coverage of HTTQ23 and HAP40 for HL2 and HD4 HDX-MS. For HTTQ23 (HAP40), 388 (58) peptides, represented by navy blue rectangles, were analyzed to generate a sequence coverage of 76.8 % (82.0 %) and a peptide-per-reside redundancy of 1.74 (2.10).

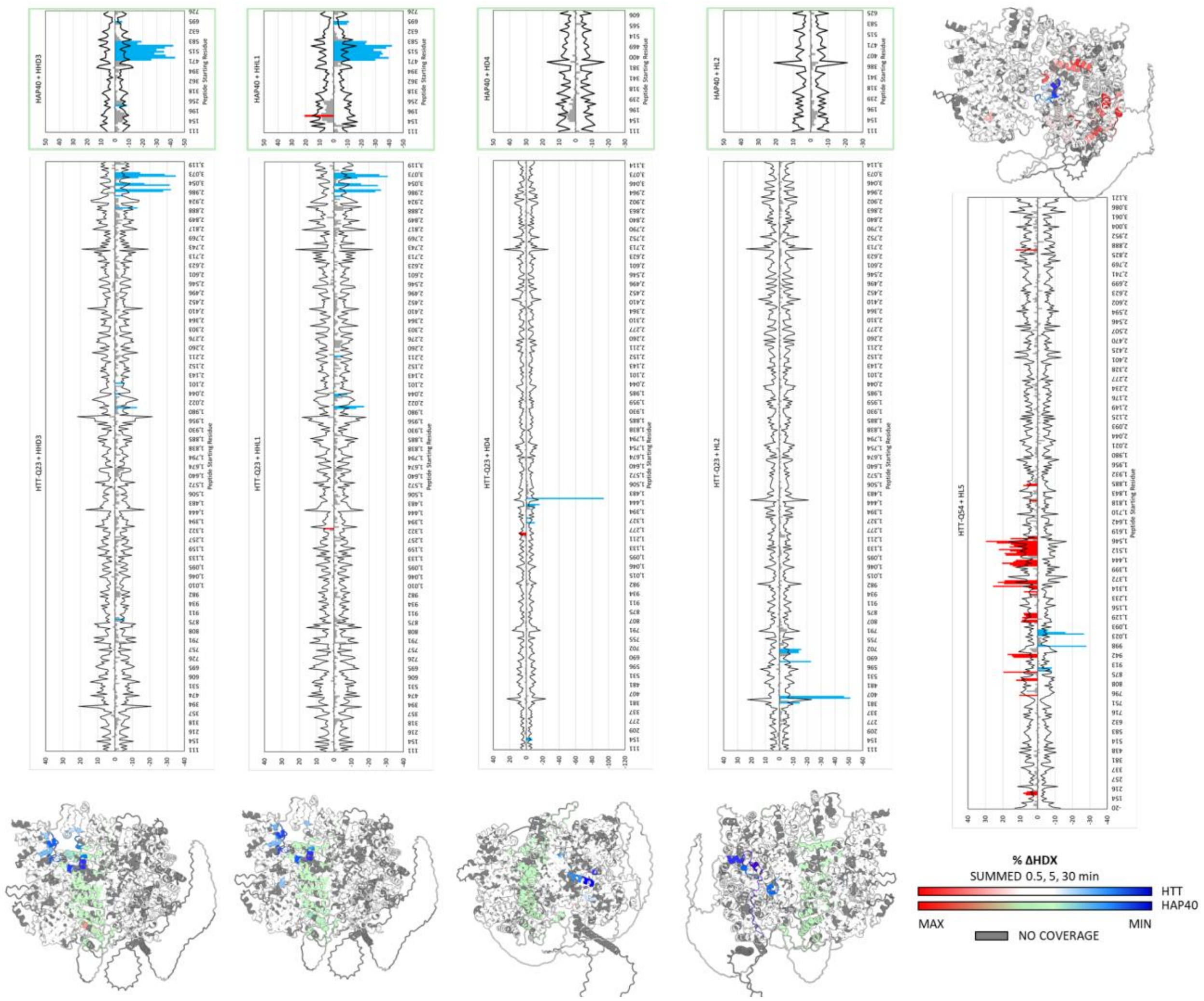


**Supplementary Figure 7.** ΔHDX-MS bar plots. For the cumulative differences to be considered statistically significant, they must exceed the cumulative error (propagated from 3 std deviations of each mean of fractional uptake at each timepoint).

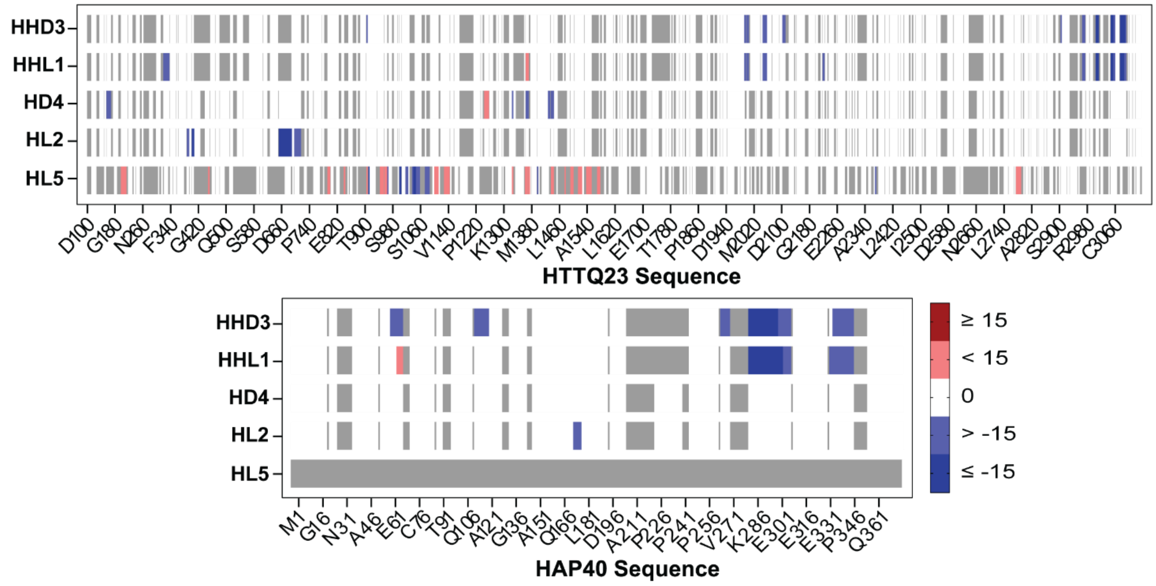


**Supplementary Figure 8.** Differential Hydrogen-Deuterium Exchange Mass Spectrometry (ΔHDX-MS) of HTTQ23-HAP40 + HHD3, HHL1, HD4, or HL2, and HTTQ23 + HL5. Cumulative differences (0.5, 5, 30 mins) in fractional uptake exceeding cumulative error are shown as increases (red), decreases (blue), statistically insignificant (white), and no coverage (grey). Per-residue heatmaps sequence numbering is based on HTTQ23 (3144 aa., NCBI Reference Sequence: NP_002102.4).

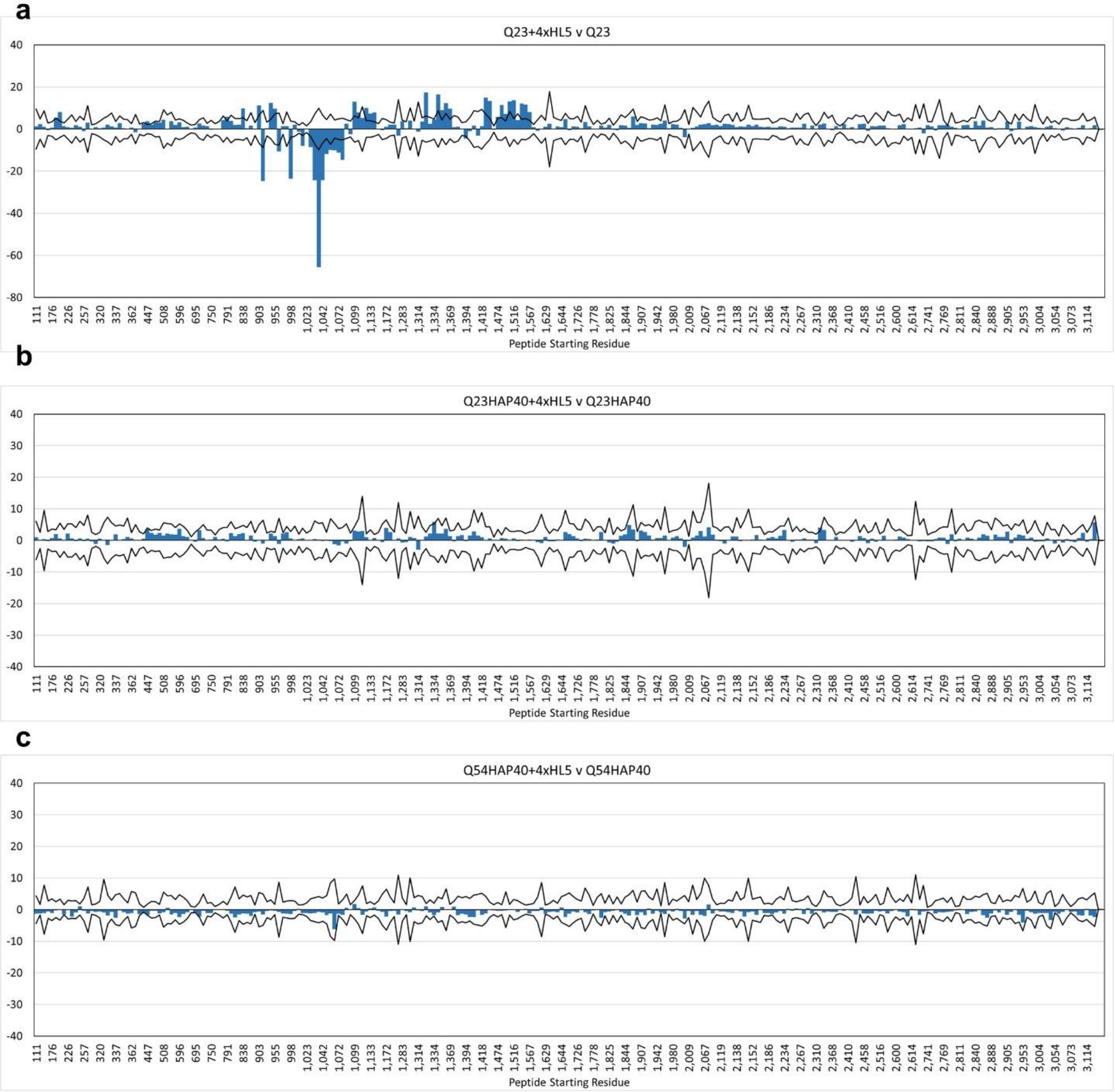


**Supplementary Figure 9.** ΔHDX-MS bar plots of HL5 impact on a, HTTQ23, b, HTTQ23-HAP40, and c, HTTQ54-HAP40. For the cumulative differences of a peptide at 3 timepoints (0.5, 5, 30 min) to be considered statistically significant, they must exceed the cumulative error (propagated from 3 sigma of each mean of fractional uptake). Though HTTQ23-HAP40 results indicate minor cumulative ΔHDX increases (<1%) in peptides SSNPSKSQGRAQRLGSSSVRPGLY (1334-1357) and LLSPERRTNTPKAISE (2328-2343), these were not observed for Q54-HAP40 and are likely just noise in the data set.

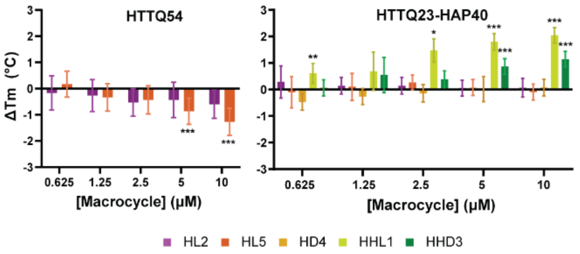


| | HTTQ54 | | HTTQ23-HAP40 | |
|---|---|---|---|---|
| | ΔTm (°C) | P value | ΔTm (°C) | P value |
| HL2 | -0.60 | 0.02 | 0.07 | 0.62 |
| HL5 | -1.27 | 0.0006 *** | 0.08 | 0.43 |
| HD4 | No data | No data | 0.10 | 0.15 |
| HHL1 | No data | No data | 2.04 | 0.00007 *** |
| HHD3 | No data | No data | 1.14 | 0.00001 *** |

**Supplementary Figure. 10.** Thermal shift ($\Delta T_m$) induced by macrocycle binding. Melting temperature of HTTQ54 (left) with HL5 or HL2 minus HTTQ54 with DMSO (46.5 °C), and (right) HTTQ23-HAP40 with HD4, HL2, HL5, HHD2, HHD3, or HHL1 minus HTTQ23-HAP40 with DMSO (51.0 °C). $\Delta T_m$ significance was determined using an unpaired t-test with Welch correction, comparing to the mean $T_m$ of Q54 or Q23-HAP40 with DMSO; *P < 0.05, **P < 0.01, ***P < 0.001. The summary table reports the $\Delta T_m$ and exact P value at the highest concentration tested (10 μM).

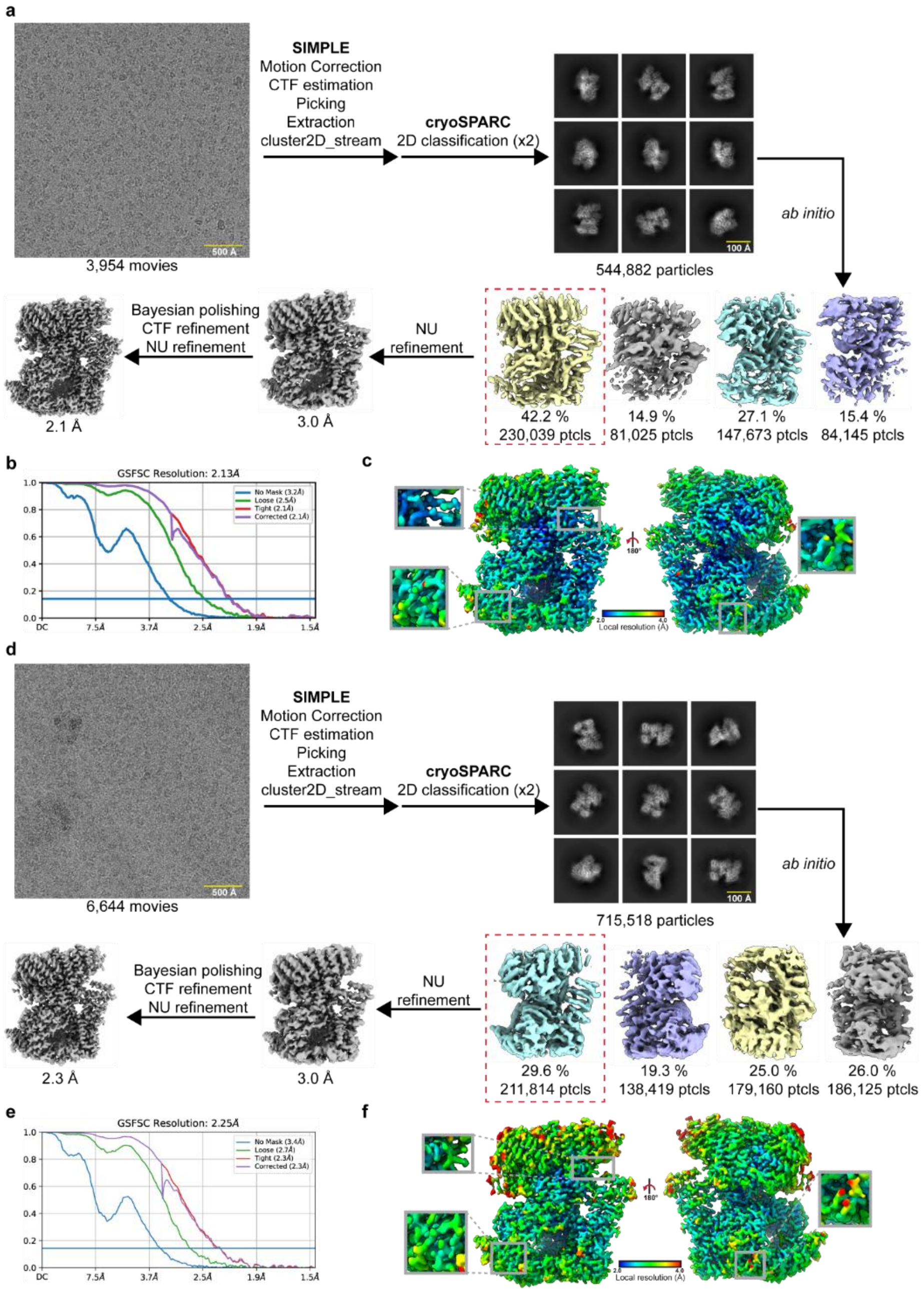


**Supplementary Figure 11.** Cryo-EM processing workflow and resolution metrics for HTT-HAP40 macrocycle complexes. ***a***, Image processing workflow for HTT-HAP40-HHL1-HL2-HD4-HHD2, with global (**b**) and local (**c**) resolution estimates. ***d***, Image processing workflow for HTT-HAP40-HHD3-HL2-HD4-HHD2, with global (**e**) and local (**f**) resolution estimates.

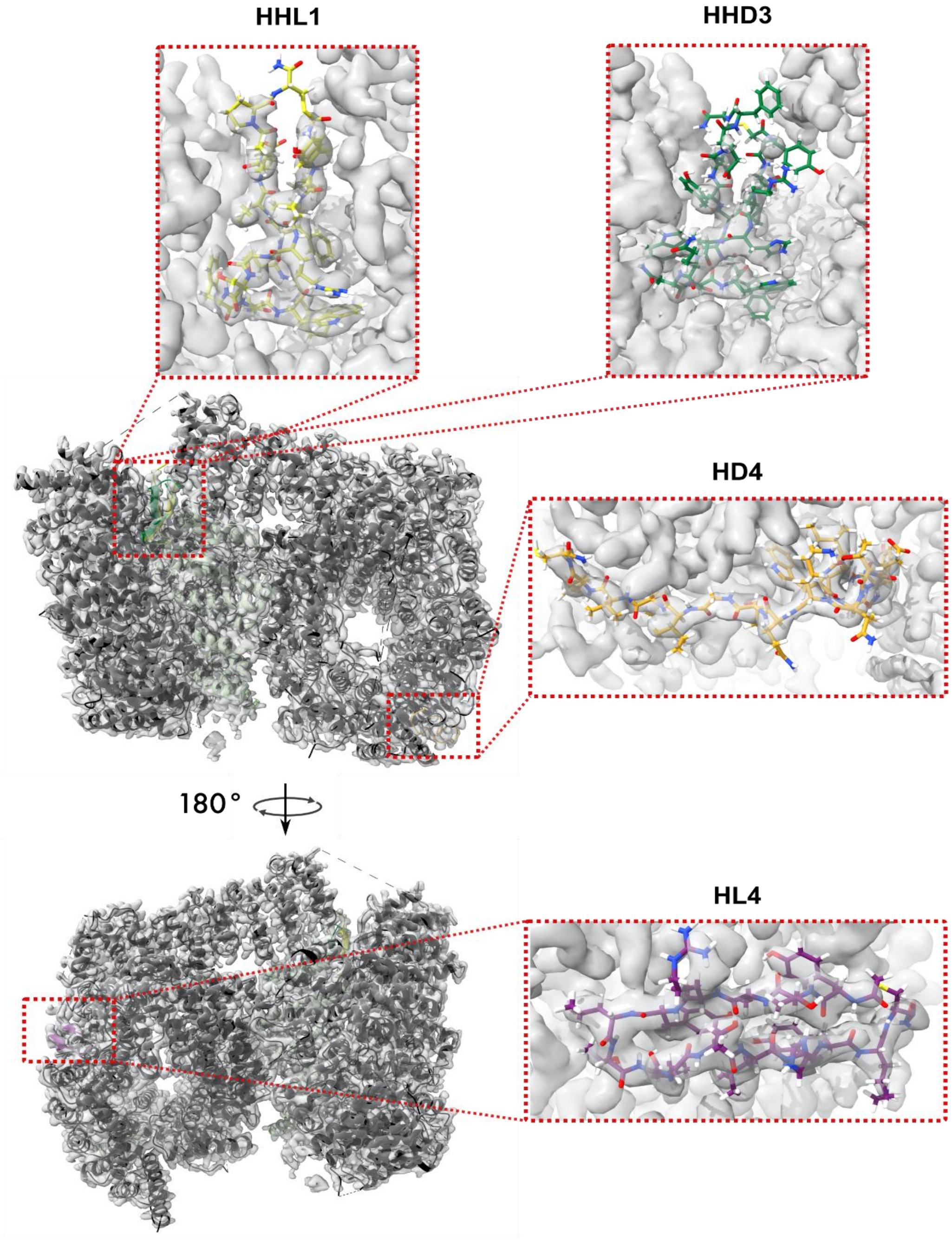


**Supplementary Figure 12.** Cryo-EM final map to model fit. The final map of HTT-HAP40 with macrocycles HHL1, HHD3, HD4 and HL2 is shown in grey, with the HTT (black), HAP40 (light green), HHL1 (dark green) HHD3 (bright green), HL2 (purple) and HD4 (yellow) models shown as ribbons. Insets show the map-model fits of the four macrocycles.

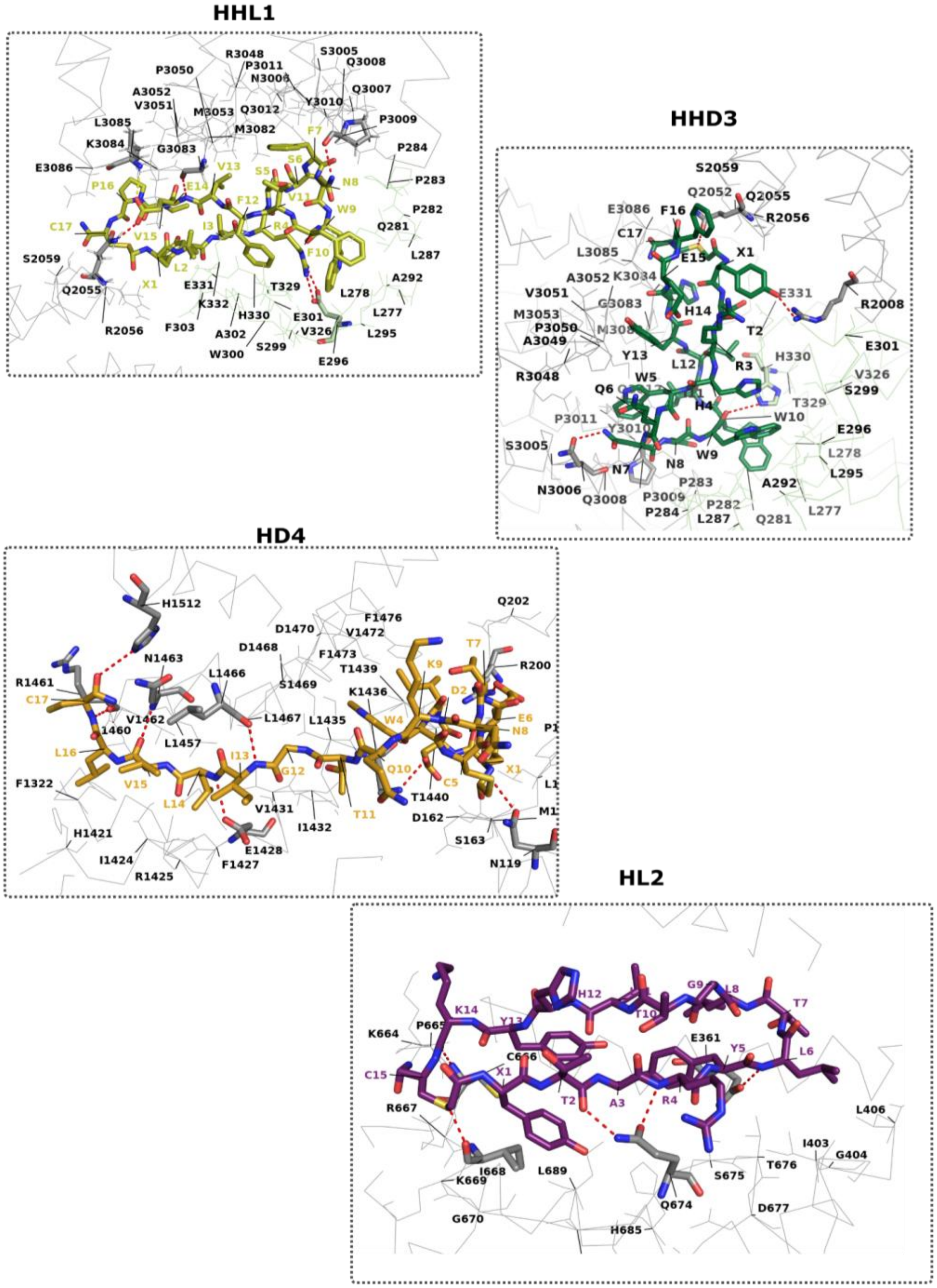


**Supplementary Figure 13.** Binding pockets of macrocycles bound to HTT-HAP40. The HTT backbone is shown as a grey ribbon, while the HAP40 backbone as a pale green ribbon. The residues in the binding pocket are shown as lines with labels. Residues forming hydrogen bonds or salt bridges are shown as sticks with the bonds shown as red dashed lines. The macrocycles are shown as sticks, with color coding as shown previously.

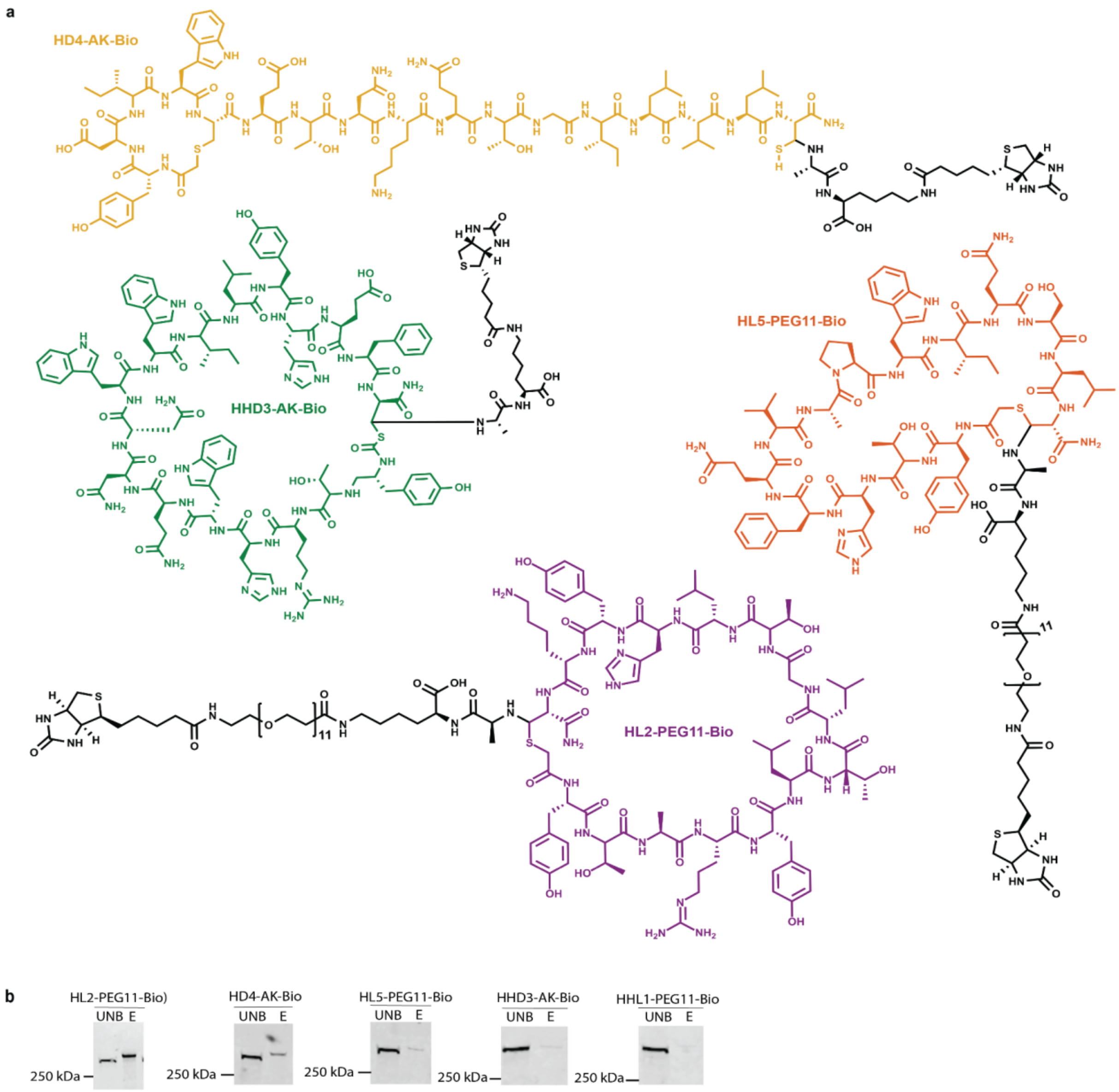


**Supplementary Figure 14. Biotinylated macrocycles.** ***a***, HD4 and HHD3 were extended with an alanine-lysine linker (AK-Bio) and HL2 and HL5 were extended via a PEG11 spacer (PEG11-Bio). ***b***, Western blot validation of HTT pulldown. Input=2%, Unbound=2% and Elution=2%. Anti-HTT antibody MA3-040

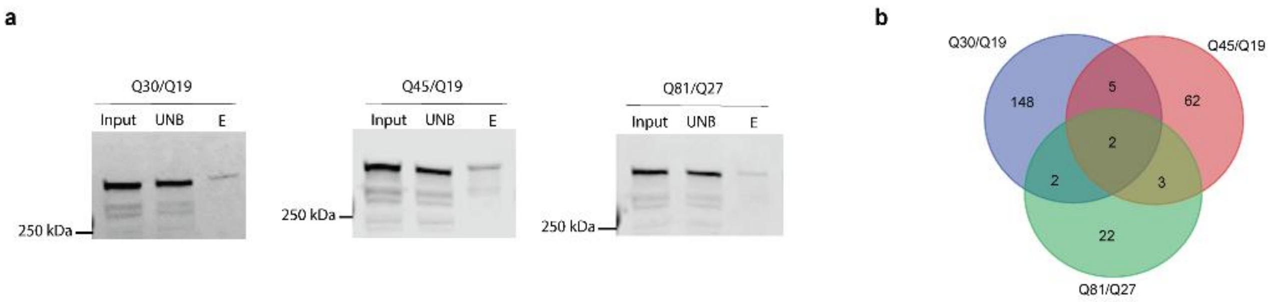


**Supplementary Figure 15.** ***a***, Western blot validation of HTT pulldown in NPCs. Input=2%, Unbound=2% and Elution=2%. Anti-HTT antibody D7F7 ***b,*** Venn diagram comparing proteins identified with >2-fold enrichment compared to control.

**Supplementary Table 1.** Cryo-EM data collection, refinement and validation statistics

| | HTT-HAP40-HHL1-HL2-HD4-HHD2 (PDB 9PMW) (EMD-71743) | HTT-HAP40-HHD3-HL2-HD4-HHD2 (PDB 9PN0) (EMD-71744) |
|---|---|---|
| **Data collection and processing** | | |
| Magnification | 165,000 | 165,000 |
| Voltage (kV) | 300 | 300 |
| Electron exposure (e–/Å$^2$) | 51.8 | 51.8 |
| Defocus range (μm) | -2.0 to -0.5 | -2.0 to -0.5 |
| Pixel size (Å) | 0.732 | 0.732 |
| Symmetry imposed | C1 | C1 |
| Initial particle images (no.) | 988,456 | 1,661,026 |
| Final particle images (no.) | 230,039 | 211,814 |
| Map resolution (Å) | 2.1 | 2.3 |
| FSC threshold | 0.143 | 0.143 |
| Map resolution range (Å) | 1.9-35 | 2.0-39 |
| **Refinement** | | |
| Initial model used (PDB code) | 6X9O | 6X9O |
| Model resolution (Å) | 2.1 | 2.3 |
| FSC threshold | 0.143 | 0.143 |
| Map sharpening *B* factor (Å$^2$) | deepEMhancer | deepEMhancer |
| Model composition | | |
| Non-hydrogen atoms | 21400 | 20419 |
| Protein residues | 2678 | 2546 |
| Ligands | 49 | 49 |
| *B* factors (Å$^2$) | | |
| Protein | 0-95 | 18-95 |
| Ligand | 0-30 | 0-30 |
| R.m.s.z[1] | | |
| Bond lengths | 0.61 | 0.72 |
| Bond angles | 1.0 | 1.06 |
| Validation | | |
| MolProbity score | 0.68 | 0.88 |
| Clashscore | 0.14 | 0.24 |
| Poor rotamers (%) | 0.76 | 1.06 |
| Ramachandran plot | | |
| Favored (%) | 97 | 96 |
| Allowed (%) | 3 | 4 |
| Disallowed (%) | 0 | 0 |

[1] Z-score for the bond length or angle, the number of standard deviations the value is removed for the expected value. Z >5 is considered an outlier.

**Supplementary Table 2.** Analyses of HTT-HAP40 interfaces upon binding macrocycles

| **Macrocycle** | **Interface area ($A^2$)** | **$*N_{SB}+N_{HB}+N_{pi}$** | **ΔG (kcal/mol)** | **Residues in binding pocket (HTT-*HAP40*)** |
|---|---|---|---|---|
| HHL1 | 1049 | 10 | -14.8 | Q2055, R2056, S2059, S3005, N3006, Q3007, Q3008, P3009, Y3010, P3011, Q3012, R3048, P3050, V3051, A3052, M3053, M3082, G3083, K3084, L3085, E3086, *L277, L278, Q281, P282,P283, P284, L287, A292, L295, E296, S299, W300, E301, A302, F303, V326, T329, H330, D331, K332* |
| HHD3 | 1076 | 4 | -13.7 | R2008, Q2052, Q2055, R2056, S2059, R2063, S3005, N3006, Q3008, P3009, Y3010, P3011, Q3012, R3048, A3049, P3050, V3051, A3052, M3053, M3082, G3083, K3084, L3085, E3086, *L277, L2278, Q281, P282, P283, P284, L287, A292, L295, E296, S299, E301, V326, H330, E331* |
| HD4 | 1021.1 | 11 | -15.1 | N119, M161, D162, S163, L165, P166, L198, R200, Q202, H1421, I1424, R1425, F1427, E1428, V1431, I1432, L1435, K1436, T1439, T1440, L1457, L1460, R1461, V1462, N1463, L1466, L1467, D1468, S1469, D1470, V1472, F1473, F1476, H1512 |
| HL2 | 755.0 | 5 | -9.1 | V302, V304, E305, E354, V357, Q358, E361, L362, L364, H365, Q368, Q370, I403, G404, L406, K664, P665, C666,R 667, I668, K669, G670, I672, G673, Q674, S675, T676, D677, H685, L689 |

*Number of salt bridges + number of hydrogen bonds + number of pi-stacking interactions.
Underlined residues participate in hydrogen bonds/salt bridges
